# Importin α7 deficiency causes infertility in male mice by disrupting spermatogenesis

**DOI:** 10.1101/2020.11.09.374652

**Authors:** Na Liu, Fatimunnisa Qadri, Hauke Busch, Stefanie Huegel, Gabin Sihn, Ilya Chuykin, Enno Hartmann, Michael Bader, Franziska Rother

## Abstract

Spermatogenesis is driven by an ordered series of events, which rely on trafficking of specific proteins between nucleus and cytoplasm. The importin α family of proteins mediates movement of specific cargo proteins when bound to importin β. Importin α genes have distinct expression patterns in mouse testis, implying they may have unique roles during mammalian spermatogenesis. Here we use a loss-of-function approach to specifically determine the role of importin α7 in spermatogenesis and male fertility. We show that ablation of importin α7 in male mice leads to infertility and has multiple cumulative effects on both germ cells and Sertoli cells. Importin α7-deficient mice exhibit an impaired Sertoli cell function, including loss of Sertoli cells and a compromised nuclear transport of the androgen receptor. Furthermore, our data demonstrate devastating defects in spermiogenesis that are accompanied by disturbed histone-protamine-exchange, absence of the transcriptional regulator Brwd1 and altered expression of Rfx2 target genes, resulting in incomplete sperm maturation and massive loss of sperms. Our work uncovers the essential role of importin α7 in spermatogenesis and hence in male fertility.

## Introduction

The best characterized mechanism of nuclear import of molecules consists of importin α and importin β heterodimers. Importin α proteins are comprised of three main structural domains: a N-terminal region which is the importin β binding (IBB) domain; a central domain containing Armadillo (ARM) motifs; and a weakly conserved C-terminal region. The central domain of importin α binds Nuclear Localization Signals (NLSs) that are present in the target cargo proteins. Upon cargo binding, importin α binds to importin β via its IBB domain forming a trimeric transport complex, which is then translocated into the nucleus via importin β interactions with nucleoporins (Nups) lining the nuclear pore complex (Macara, 2001; Miyamoto et al., 2012). To date, three importin α subtypes have been identified in *C. elegans* and *D. melanogaster*, while up to seven importin α isoforms have been found in mammals (Kohler et al., 1997; Tejomurtula et al., 2009; Tsuji et al., 1997).

Male reproductive function relies on normal spermatogenesis within the seminiferous epithelium of the testis. During spermatogenesis, the spermatogonia undergo mitosis and differentiate into primary spermatocytes. The primary spermatocytes process through preleptotene, leptotene, zygotene, pachytene, and diplotene stages of meiosis I to generate secondary spermatocytes. Subsequently, the secondary spermatocytes enter the second meiotic division resulting in round spermatids. The haploid round spermatids undergo dramatic morphological changes, and finally differentiate into mature spermatozoa (Russell, 1990).

Sertoli cells are the supporting somatic cells essential for the development of male germ cells of all stages. It has been shown that the number and function of Sertoli cells determine testicular size, germ cell numbers and spermatozoa output (Orth et al., 1988). Sertoli cell functions include (i) providing structural support and nutrition to developing germ cells, (Chihara et al., 2013) simultaneous support of synchronous differentiation among several cohorts of germ cells of differing maturity, (iii) secretion of seminiferous fluid, (iv) phagocytosis of degenerating germ cells and residual bodies and (v) release of spermatids at spermiation (Bellve and Zheng, 1989; Chihara et al., 2013; Clermont, 1993; Russell and Griswold, 1993).

A key feature of Sertoli cell structural support for developing germ cells is the blood-testis barrier (BTB) that resides in tight junctions located between adjacent Sertoli cells (Johnson et al., 2008). At the beginning of meiosis preleptotene spermatocytes ‛pass through’ the BTB. Once the germ cells move beyond the BTB and the BTB reforms behind them, the germ cells no longer have access to serum factors and they become totally dependent upon Sertoli cells to supply nutrients and growth factors (Walker, 2010). This structural arrangement creates an immunologic barrier by isolating the more advanced germ cell types from the immune system so that their antigens do not stimulate autoimmunity (Johnson et al., 2008; Orth et al., 1988). The successful completion of spermatogenesis is dependent on successive division and differentiation steps, which require multiple changes in gene expression; coordinated by a plethora of transcription and other factors expressed within the testis (Eddy and O’Brien, 1998; Hermo et al., 2010). The access of these factors to the nucleus where they can exert their function, is tightly regulated for these proteins, and it has been postulated that germ cell differentiation is controlled by nucleocytoplasmic transport events (Major et al., 2011). In fact, the mRNAs of different importin α isoforms and of importin β are all expressed in spermatogonia, spermatocytes, round spermatids and Sertoli cells (Major et al., 2011; Shima et al., 2004), which raises the possibility that the importin α/β mediated nuclear import pathway is involved in the regulation of spermatogenesis and Sertoli cell function. However, a specific role of a single importin α isoform in spermatogenesis and male reproduction has not been determined yet.

We have previously shown that in importin α7-deficient mothers, embryonic development stops at the two-cell stage due to a severely disturbed zygotic genome activation, therefore importin α7 is essential for early embryonic development in mice (Rother et al., 2011). In this report, we show that ablation of Kpna6, the gene encoding for importin α7, results in a critical defect in spermatogenesis in male mice. We demonstrate that importin α7 protein is expressed in the nuclei of round spermatids, elongating spermatids and Sertoli cells. Consistent with this pattern, importin α7 deficiency results in multiple defects in both germ cells and Sertoli cells culminating in oligozoospermia. Our results demonstrate an essential role for importin α7 in male fertility by regulating spermiogenesis and Sertoli cell function.

## Results

### Importin α7 is essential for male fertility

To investigate the physiological role of mammalian importin α7, we have generated two mouse lines with targeted disruption of importin α7. In one line, exon 2 of the importin α7 gene is deleted which, due to unexpected alternative splicing, results in a shortened mRNA containing a cryptic translational start site in exon 3 and thus leading to synthesis of a truncated protein lacking the importin ß binding domain (α7^ΔIBB/ΔIBB^, Fig. 1A). In the other line a gene trap cassette is located in intron 1 of the importin α7 gene resulting in a complete loss of the protein (α7^−/−^, Fig. 1A). Female mice of both lines are infertile (Rother et al., 2011). Most interestingly, the male α7^−/−^ mice are fertile, while α7^ΔIBB/ΔIBB^ males were found to be sterile, although they were sexually active and produced vaginal plugs in female partners (data not shown). We observed that, although importin α7 protein is disrupted in all other organs of α7^−/−^ males (Rother et al., 2011), full-length importin α7 protein is still expressed in the testis, albeit to a lower extent, whereas it is completely absent from α7^ΔIBB/ΔIBB^ testes (Fig. 1B). The reason for the exclusive expression of importin α7 in the testis of importin α7^−/−^ males is that an alternative promoter and exon 1 (exon 1A) are used which are located downstream of the gene trap cassette (Fig. 1A-C, EST accession number BY353738.1). On the contrary, only a truncated non-functional importin α7 is existent in the α7^ΔIBB/ΔIBB^ testes (Fig. 1B), which leads to male infertility, suggesting that importin α7 plays an important role in mouse testis, and is essential for male fertility.

**Fig. 1.**
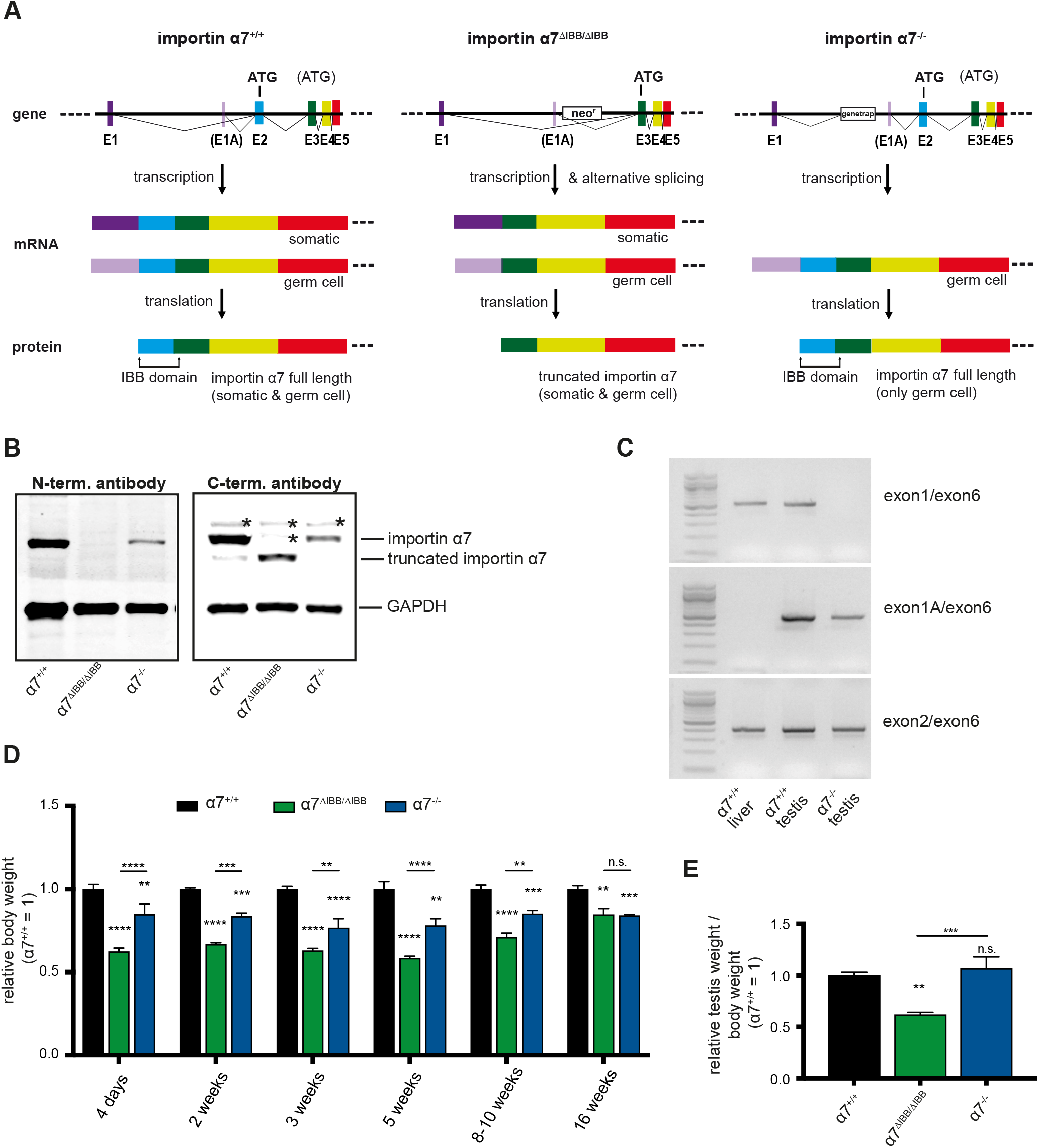
Disruption of importin α7 causes growth retardation and reduced testis size. (A) Gene targeting strategy for importin α7^ΔIBB/ΔIBB^ and α7^−/−^ mice. In α7^ΔIBB/ΔIBB^ mice, exon 2 is replaced by a neomycin resistence (neo^r^) cassette with a polyadenylation site (pA). Since transcription does not always stop at pA, a splicing variant is generated, carrying an *in frame* translational start site in exon 3. This results in the formation of a truncated protein lacking the IBB. In importin α7^−/−^ mice, a genetrap is located in intron 1, resulting in complete loss of the protein in most of the tissues. However, a testis-specific exon 1A (E1A) allows the generation of a germ cell-specific mRNA which in turn results in the generation of a full-length protein. (B) Western blot analysis of importin α7 expression in the testes of WT (α7^+/+^), α7^ΔIBB/ΔIBB^ and α7^−/−^ mice. The 58 kDa full-length importin α7 protein is absent in the α7^ΔIBB/ΔIBB^ testis, while it can be detected in both α7^−/−^ and WT testes (left panel). An importin α7^ΔIBB^ protein which is about 10 kDa smaller than the full-length importin α7 protein is found in the testis of α7^ΔIBB/ΔIBB^ mice (right panel). The antibody against the N-terminus recognizes the full-length (α7^+/+^, α7^−/−^), but not the truncated (α7^ΔIBB/ΔIBB^) importin α7. The asterisks mark unspecific cross-reactions of the antibody. (C) RT-PCR of WT liver, WT testis and importin α7^−/−^ testis using primer pairs spanning different exons. In WT liver and testis, a transcript spanning exon1 and exon6 can be detected, while it is absent in α7^−/−^ testis. Only WT testis, but not WT liver expresses a specific transcript using an alternative exon (exon1A) and this transcript can be detected in α7^−/−^ mice as well, albeit to a lesser extent than in WT. As a control, all tested tissues express importin α7 transcripts spanning exon2 and exon6. (D) Relative body weight of WT, importin α7^ΔIBB/ΔIBB^ and α7^−/−^ mice at 4 days, and 2, 3, 5, 8-10, 15-16 weeks of age (α7^+/+^=1, n=6 per group). (E) Relative testis weight compared to body weight of adult WT (α7^+/+^), importin α7^ΔIBB/ΔIBB^ and α7^−/−^ mice at 8-10 weeks of age (n=6 per group).

### Disruption of importin α7 gene causes growth retardation, reduced testis size and severe oligozoospermia

Importin α7^ΔIBB/ΔIBB^ mice and α7^−/−^ mice were born at a lower frequency (α7^ΔIBB/ΔIBB^: 18.8%, n=739; p<0.0001; α7^−/−^: 16%, n=214, p=0.0015) than predicted by Mendelian laws. With regards to growth and development, the heterozygous males are indistinguishable from wildtype (WT) males (data not shown). However, α7^ΔIBB/ΔIBB^ pups that survived the gestation period displayed severe growth retardation in the postnatal phase, and this growth defect persisted until adult life (Fig. 1D). Male importin α7^−/−^ mice also displayed a significant growth retardation, albeit the effect in young mice was stronger in α7^ΔIBB/ΔIBB^ mice. At the age of 16 weeks, males of both mutant lines showed the same reduction in body weight compared to WT males (Fig. 1D). Testes of adult α7^ΔIBB/ΔIBB^ mice exhibited a pronounced reduction both in size and weight, and the testicular weight to body weight ratio was reduced by 40% at the age of 8-10 weeks compared to WT controls and to α7^−/−^ mice, which displayed a normal relative testis weight and size (Fig. 1E). Plasma testosterone levels were unchanged (data not shown). Histological analyses of testes revealed that spermatogenesis was drastically altered in importin α7^ΔIBB/ΔIBB^ while no major changes were detected in importin α7^−/−^ compared to WT testes (Fig. 2A). Seminiferous tubules in the importin α7^ΔIBB/ΔIBB^ testes were smaller in diameter than those in WT and importin α7^−/−^ mice (Fig. 2A and C). The number of germ cells was reduced, and they were disorganized in the tubular epithelium. Moreover, mature spermatozoa were rarely found in the lumen of α7^ΔIBB/ΔIBB^ seminiferous tubules and multinucleated spermatid giant cells were frequently observed (Fig. 2A). There were very few spermatozoa in the caput of the α7^ΔIBB/ΔIBB^ epididymides, and spermatozoa were hardly detectable in the caudal epididymides by H&E staining (Fig. 2B). Additionally, sloughed germ cells, and germ cell debris were commonly observed in the corpus epididymal lumen of α7^ΔIBB/ΔIBB^ males (Fig. 2B). The total epididymal sperm number in α7^ΔIBB/ΔIBB^ was only 3% of those of WT males (Fig. 2D; 0.8×10^6^ versus 25.6×10^6^), moreover, almost all of the residual sperms found in the α7^ΔIBB/ΔIBB^ epididymides displayed abnormal heads in contrast to α7^−/−^ and WT sperms (Fig. 2E). Surprisingly, also importin α7^−/−^ sperm count was significantly reduced, suggesting a partially reduced fertility in these mice (Fig. 2D). In both lines the epididymal sperm count of heterozygous mice was normal (Fig. S1A).

**Fig. 2.**
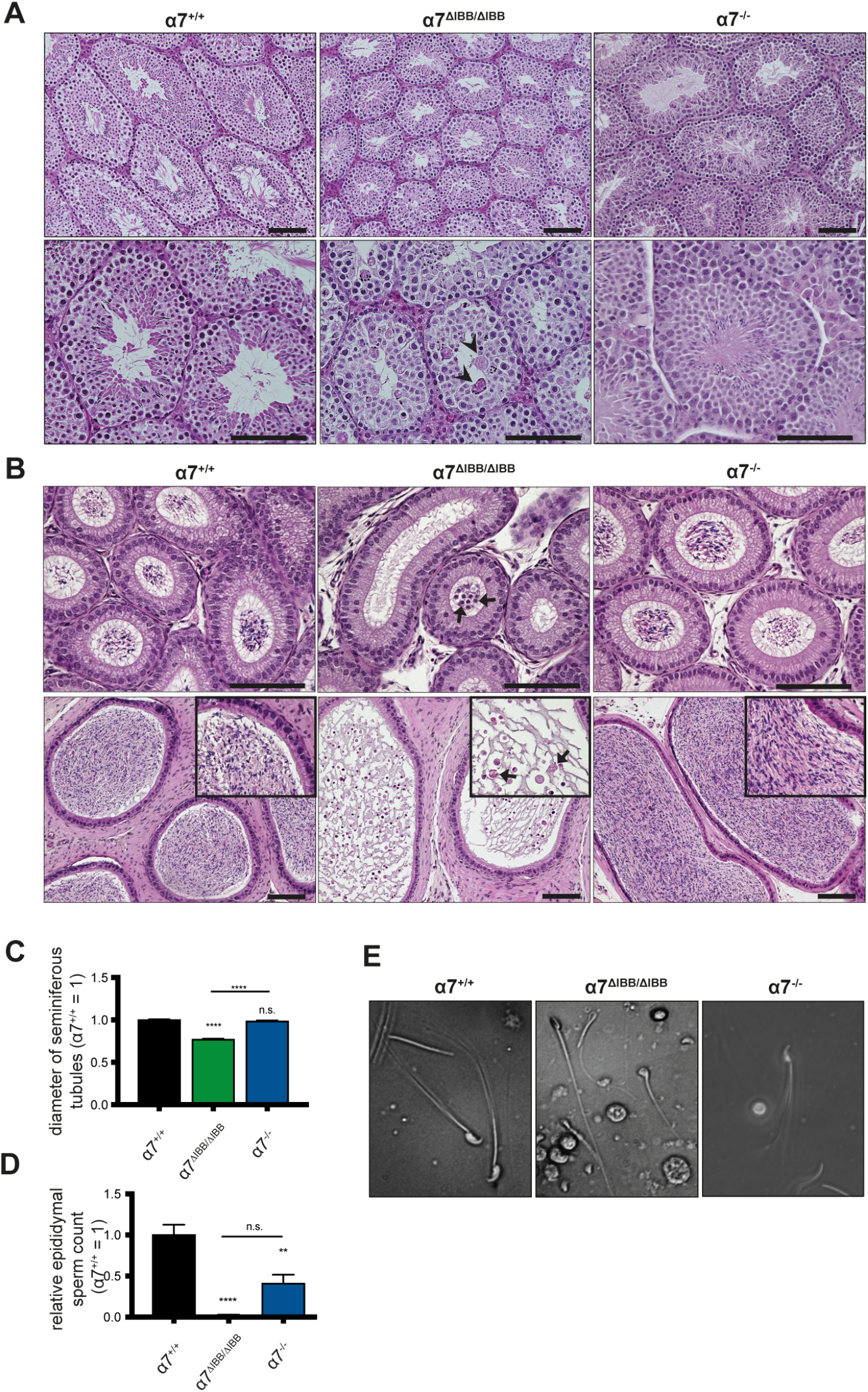
Disruption of importin α7 causes oligozoospermia. (A) H&E staining of paraffin embedded testis sections of adult WT (α7^+/+^), importin α7^ΔIBB/ΔIBB^ and α7^−/−^ mice (upper lane 20x; lower lane 40x magnification). (B) H&E staining of epididymis sections of adult WT (α7^+/+^), importin α7^ΔIBB/ΔIBB^ and α7^−/−^ mice. Upper lane: caput epididymidis (40x); lower lane: cauda epididymidis (20x, inserts 40x). Arrow heads in (A): multinucleated spermatid giant cells; arrows in (B): immature germ cells (upper lane), sloughed immature germ cells (lower lane). (C) Relative diameters of seminiferous tubules in importin α7^ΔIBB/ΔIBB^ and α7^−/−^ mice compared to WT (α7^+/+^). (D) Epididymal sperm count of WT (α7^+/+^), importin α7^ΔIBB/ΔIBB^ and α7^−/−^ mice (n=4-6). (E) Representative images of epididymal sperms from WT (α7^+/+^), importin α7^ΔIBB/ΔIBB^ and α7^−/−^ mice. Scale bars: 100 μm.

### Importin α7 expression pattern in mouse testis

To assess the cell type-specific function of importin α7 in the testis, we performed immunohistochemistry for importin α7 in WT mice using an antibody which detects the C-terminus and thereby the full length as well as the truncated form of importin α7 (Fig. 3). No importin α7 protein could be detected in spermatogonia and meiotic spermatocytes. Early round spermatids (step 1) showed very low levels of expression which increased throughout the development of round spermatids (stages I-VIII), reaching its highest expression in step 9 elongating spermatids (stage IX), where importin α7 displayed a high nuclear and low cytoplasmic expression. With the onset of nuclear elongation, localization of importin α7 shifted to the cytoplasm, and was no longer detectable after the residual bodies were removed in step 16 sperms (stage VII-VIII). Importin α7 was not detected in peritubular myoid cells and displayed a low nuclear expression in some of the Leydig cells, however it is highly expressed in the nuclei of Sertoli cells in all stages of the seminiferous epithelium (Fig. 3). The massive increase of importin α7 expression in step 9 elongating spermatids, and the high expression level in the nuclei of Sertoli cells suggest an important role of importin α7 in these cells. As the antibody against the C-terminus of importin α7 also detects the truncated form of the protein, it showed a regular staining in α7^ΔIBB/ΔIBB^ testes, while testis sections of importin α7^−/−^ mice that express importin α7 exclusively in the testes revealed that the protein expression is rescued in germ cells, but not in Sertoli cells in this mouse line (Fig. 4A). To verify these results, we generated an antibody against the N-terminus of importin α7 that could discriminate between the full-length and the truncated ΔIBB-protein in which the N-terminus is missing (Fig. 1A). Staining of testis sections of WT mice showed a robust signal in round and elongating spermatids as well as Sertoli cells. In contrast, no signals could be detected in α7^ΔIBB/ΔIBB^ testis sections with this antibody, confirming the truncation of importin α7 in these mice (Fig. 4B). In importin α7^−/−^ testes, the rescued expression in developing spermatids could be verified, while no expression was found in Sertoli cells (Fig. 4B).

**Fig. 3.**
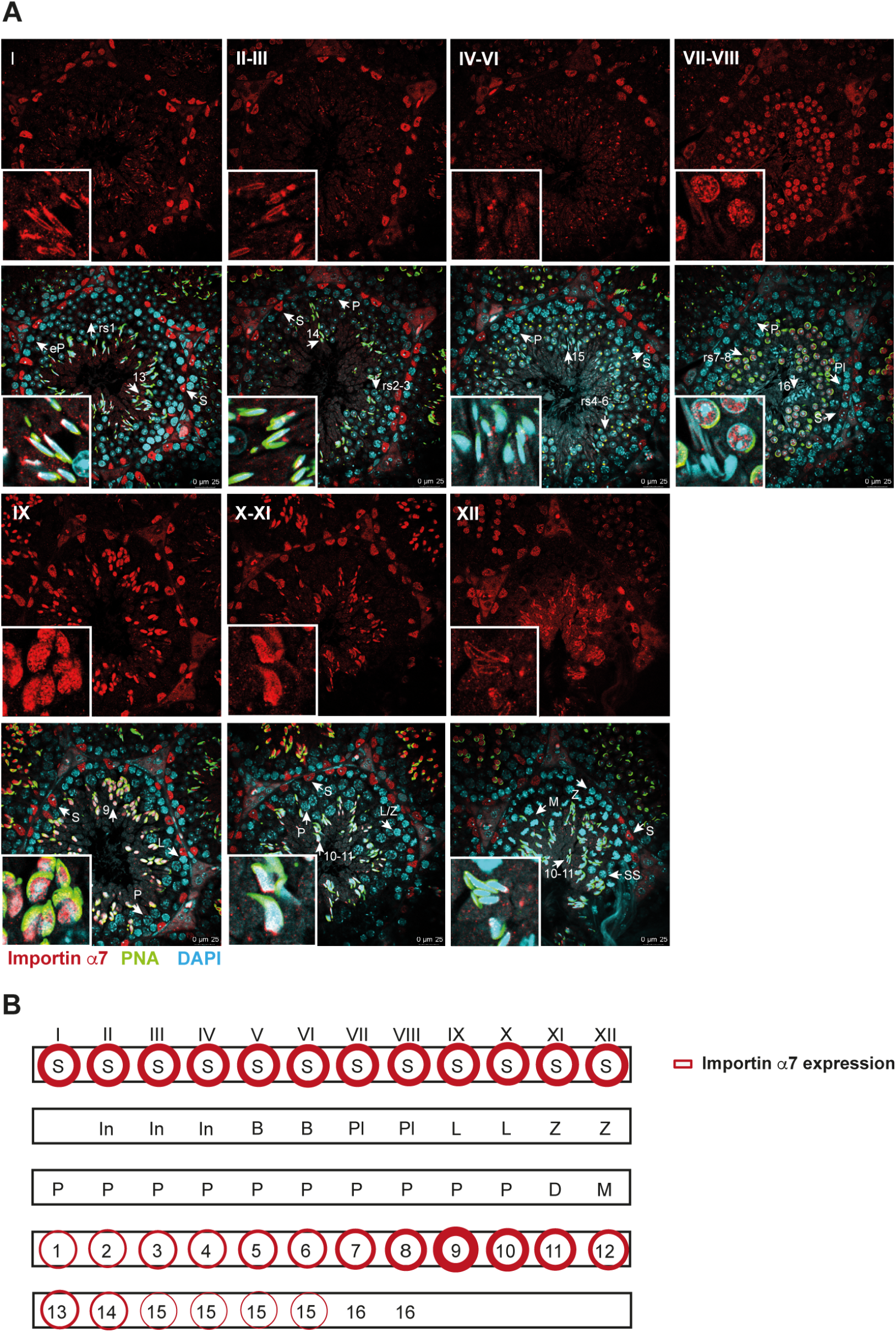
Importin α7 is expressed in postmeiotic spermatids and Sertoli cells in WT testis. (A) Immunohistochemistry of paraffin embedded WT testis sections using an antibody against the C-terminus of importin α7 (red), counterstained with DAPI (light blue) and peanut agglutinin (PNA), labelling the acrosome (green); 40x magnification. Scale bars: 25 μm. (B) Schematic image of importin α7 expressing cell types in mouse testis. S: nucleus of a Sertoli cell; eP: early pachytene spermatocyte; rs1: step 1 round spermatid; 13: step 13 sperm; P: pachytene spermatocyte; rs2-3: step 2-3 round spermatid; 14: step 14 sperm; rs4-6: step 4-6 round spermatid; 15: step 15 sperm; rs7-8: step 7-8 round spermatid; 16: step 16 sperm; Pl: preleptotene spermatocyte; L: leptotene spermatocyte; 9. Step 9 elongating spermatid; L/Z spermatocytes at leptotene/zygotene transition; Z: zygotene spermatocyte; 10-11: step 10-11 elongating spermatid; 12: step 12 sperm; M: meiosis, SS: secondary spermatocyte; D: diakinesis spermatocyte; In: intermediate spermatogonium; B: type B spermatogonium.

**Fig. 4.**
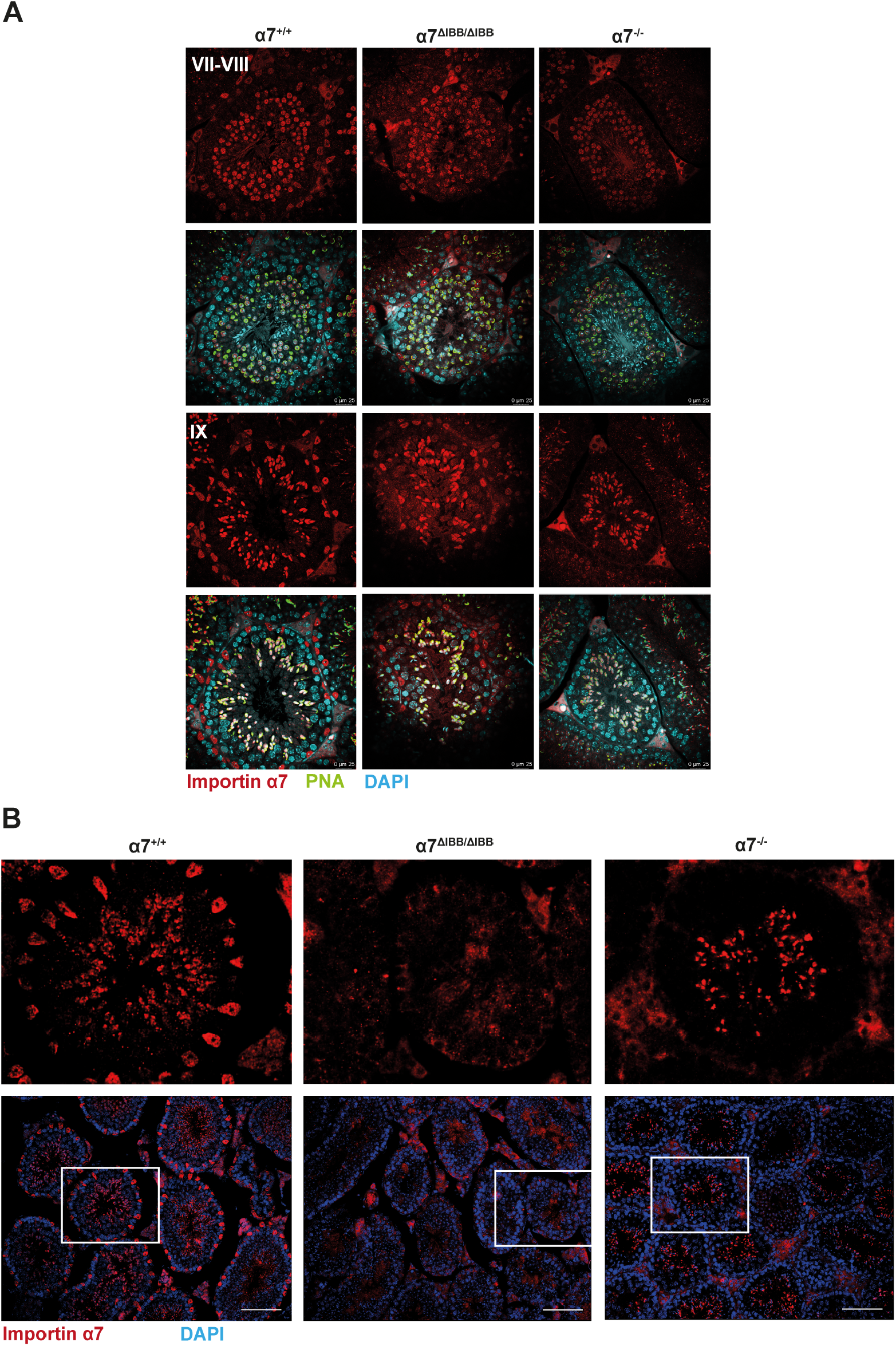
Importin α7 is expressed in developing germ cells, but not in Sertoli cells of importin α7^−/−^ mice. (A) Immunohistochemistry of WT, importin α7^ΔIBB/ΔIBB^ and α7^−/−^ testis paraffin sections using an antibody against the C-terminus of importin α7 showing selected stages of seminiferous tubules; importin α7 (red); DAPI (light blue); PNA (green); 40x magnification. (B) Immunohistochemistry of WT, importin α7^ΔIBB/ΔIBB^ and α7^−/−^ testis paraffin sections using an antibody against the N-terminus of importin α7 (red); counterstained with DAPI (blue). The protein is expressed in round and elongating spermatids, but not in Sertoli cells of importin α7^−/−^ mice, while WT mice show a robust staining in Sertoli cells, indicated in the windows. Scale bars: 100 μm.

A detailed analysis of the localization of the rescued importin α7 in importin α7^−/−^ testes using the C-terminal antibody revealed that the expression of the protein in germ cells is comparable with WT testis, with low levels in round spermatids step 1 (stage I) and gradual increase up to step 8 (stage VIII) reaching the highest level in step 9-11 elongating spermatids. In condensing spermatids (step 12) localization of importin α7 shifted to the cytoplasm as it was the case in WT mice, and was no longer detectable after the residual bodies were removed in step 16 spermatids (Fig. 4A, data not shown). We also confirmed the earlier observation that Sertoli cells of α7^−/−^ testes do not express importin α7 protein, suggesting that the expression of the protein is solely rescued in germ cells. The missing expression of importin α7 in Sertoli cells could thus account for the reduced sperm cell number observed in importin α7^−/−^ mice, however, only importin α7^ΔIBB/ΔIBB^ mice are infertile, suggesting, that the expression of the protein in germ cells is indispensable for normal sperm development and fertility.

Thus, importin α7^−/−^ mice express a mild Sertoli cell related phenotype, while importin α7^ΔIBB/ΔIBB^ mice express a mixed phenotype consisting of Sertoli cell related and germ cell related defects. The infertile importin α7^ΔIBB/ΔIBB^ mice express a truncated importin α7 protein in Sertoli cells and in the developing sperms. On the other hand, the heterozygous importin α7^ΔIBB/+^ mice express the full-length plus the truncated protein in Sertoli cells and in the developing sperms, however, sperm count in these mice turned out to be completely normal (Fig. S1A), excluding a dominant negative effect of the truncated protein on sperm count. Nevertheless, to discriminate between Sertoli cell-related phenotype and sperm cell-related phenotype, we compared mice of both lines (α7^ΔIBB/ΔIBB^ mice vs. α7^−/−^ mice). To rescue the germ cell phenotype without rescuing the Sertoli cell phenotype in importin α7^ΔIBB/ΔIBB^ mice, we crossed importin α7^ΔIBB/ΔIBB^ and α7^−/−^ mice. The resulting compound heterozygous importin α7^ΔIBB/−^ mice express only the truncated importin α7 in Sertoli cells and truncated plus full-length importin α7 in developing sperms (Fig. S1B). Epididymal sperm count revealed a significant increase of sperm number in importin α7^ΔIBB/−^ mice compared to importin α7^ΔIBB/ΔIBB^ mice (Fig. S1A), suggesting that, indeed, the addition of full-length importin α7 in developing sperms can partially rescue the oligozoospermia. However, the sperm count was still markedly lower than in WT mice and the values were comparable to importin α7^−/−^ mice, confirming that the absence of full-length importin α7 in Sertoli cells is the reason for the partial reduction in sperm count. Thus, we can conclude that importin α7 expression in spermatocytes and round spermatids is essential for normal sperm development and fertility.

### Importin α7 deficiency leads to defects in Sertoli cells

Since importin α7 was intensively expressed in the nuclei of WT Sertoli cells (Fig. 3), the protein may be essential for the function of these cells. We observed a reduced number of Sertoli cells in testes of adult α7^ΔIBB/ΔIBB^ mice and α7^−/−^ mice, suggesting that importin α7 perturbation caused a loss of Sertoli cells (Fig. 5A, wt=14.54/tubule; α7^ΔIBB/ΔIBB^=11.89/tubule; α7^−/−^ =11.39/tubule). Moreover, in both mutant lines Sertoli cells were frequently observed being detached in the middle of seminiferous tubules and the epididymal lumen (Fig. 5B).

**Fig. 5.**
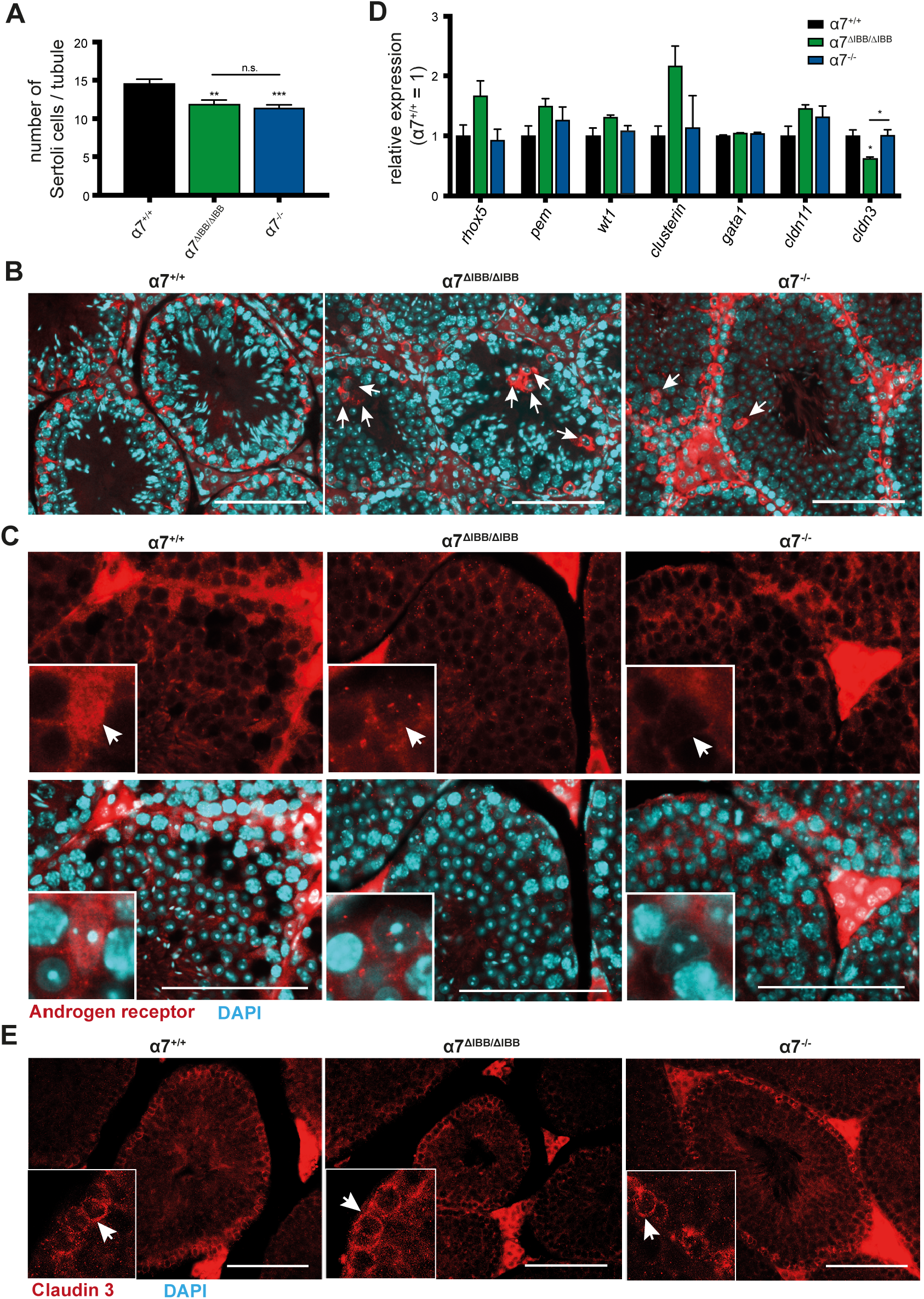
Importin α7 deficiency decreases number of Sertoli cells and prevents nuclear import of AR. (A) Number of Sertoli cells per tubule of adult WT (α7^+/+^), importin α7^ΔIBB/ΔIBB^ and α7^−/−^ testes. (B) Aberrant localization of Sertoli cells in seminiferous tubules of importin α7^ΔIBB/ΔIBB^ and α7^−/−^ testes. (C) Immunofluorescence staining of testis section against AR (red), counterstain with DAPI (light blue). Arrows mark the typical nuclei of Sertoli cells with positive signal of AR; note the nuclear localization of AR in WT and the empty nuclei in importin α7^ΔIBB/ΔIBB^ and α7^−/−^. (D) Relative expression levels of selected genes in testes of adult WT (α7^+/+^), importin α7^ΔIBB/ΔIBB^ and α7^−/−^ testes (n=6). (E) Cldn3 (red) and DAPI (blue) staining on paraffin sections of adult WT (α7^+/+^), importin α7^ΔIBB/ΔIBB^ and α7^−/−^ testes. Arrows mark Cldn3 staining of stage VIII seminiferous tubuli. Scale bars: 100 μm.

The androgen receptor (AR), which is highly expressed in Sertoli cells, plays an important role in spermatogenesis. By immunohistochemistry we observed pronounced defects of AR nuclear import in Sertoli cells of both mutant lines (Fig. 5C). However, quantification of AR mRNA expression as well as protein levels in testes did not reveal any differences (data not shown). Thus, ablation of importin α7 in Sertoli cells is accompanied by AR nuclear import deficiency. To test, whether expression of AR-related or Sertoli cell-specific genes is affected in importin α7^ΔIBB/ΔIBB^ and importin α7^−/−^ mice, we analyzed the expression levels of AR-regulated genes such as *rhox5*, *pem*, *wt1*, *clusterin*, *gata1, cldn3*, and *cldn11*. No significant differences in mRNA levels were detected for most of these genes. Only *cldn3* expression was markedly downregulated in importin α7^ΔIBB/ΔIBB^, but not in α7^−/−^ testes (Fig. 5D). To confirm these results, immunostaining of testis sections for Cldn3 was performed, revealing the specific localization in basal tight junctions of late stage VIII tubuli in both mutant mouse lines (Fig. 5E). Tight junctions are a major component of the blood-testis-barrier (BTB) located between adjacent Sertoli cells. To further analyze possible defects in tight junction formation of Sertoli cells, we performed immunostaining for ZO-1, a specific tight junction protein, in testis sections. No differences were found in the expression and localization of ZO-1 in basal tight junctions and apical ectoplasmic specializations of stage IV-VI tubules (Fig. 6A). We evaluated the functional integrity of the BTB by incubating testicular protein extracts from two months old WT mice with sera taken from either WT, importin α7^ΔIBB/ΔIBB^ or importin α7^−/−^ mice at 8, 16 and 20 weeks of age. A subsequent western blot analysis revealed in some, but not in all cases, differences in the protein band pattern with additional bands appearing in importin α7^ΔIBB/ΔIBB^ and importin α7^−/−^ mice. Thus, the immunological barrier may be leaky and therefore antibodies against testicular antigens are occasionally present in both mutant lines (Fig. 6B). However, a subsequent analysis of the presence of immunoglobulins within testicular tissue in mice of different ages revealed no differences in all three tested lines (Fig. 6C). These data suggest that the BTB is not severely impaired. Moreover, by injection of biotin into the testis we could not find a compromised BTB in importin α7^ΔIBB/ΔIBB^ and importin α7^−/−^ mice (Fig. 6D).

**Fig. 6.**
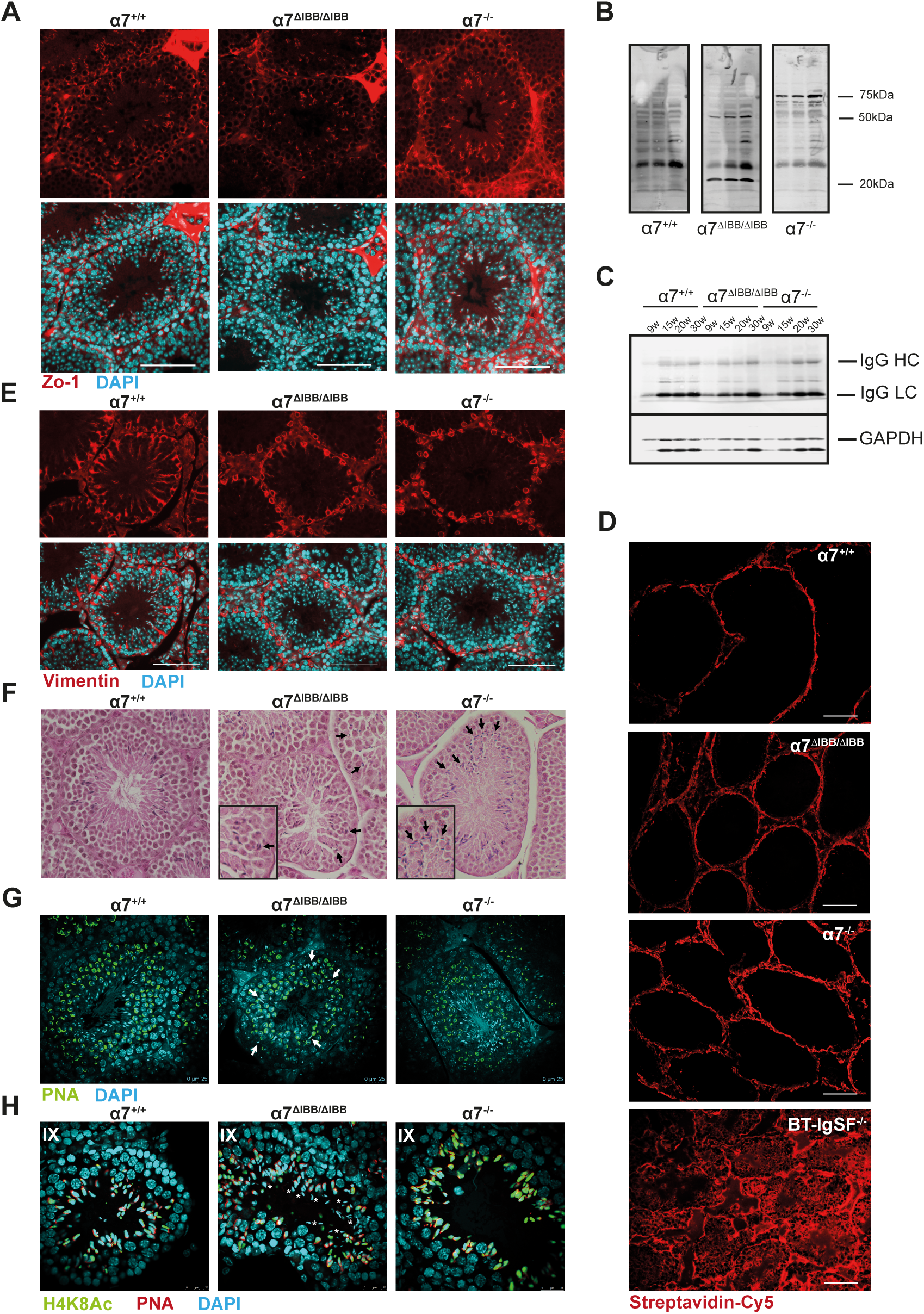
Compromised BTB and Sertoli cell function in importin α7-deficient testis. (A) ZO-1 (red) and DAPI (light blue) staining of testis paraffin sections of WT (α7^+/+^), importin α7^ΔIBB/ΔIBB^ and α7^−/−^ mice. Scale bars: 100 μm. (B) Western blot analysis of protein extracts prepared from the testes of WT mice probed with sera from WT (α7^+/+^), importin α7^ΔIBB/ΔIBB^ and α7^−/−^ mice at 16 weeks of age. Lines mark 20-, 50- and 75-kDa testicular antigens recognized by antibodies (autoantibodies) present in serum samples of different importin α7^ΔIBB/ΔIBB^ and α7^−/−^ but not WT males. (C) Representative western blot of immunoglobulins in testis extracts of importin α7^+/+^, α7^ΔIBB/ΔIBB^ and α7^−/−^ mice of different ages. (D) Biotin diffusion assay to test the integrity of the BTB. Biotin is visualized by streptavidin-Cy5 staining (red). Importin α7^ΔIBB/ΔIBB^ and α7^−/−^ tubules show no major changes of biotin distribution compared to importin α7^+/+^ seminiferous tubules. As a control, a BT-IgSF knockout mouse was analyzed, which has recently been published to show a severe disruption of the BTB (Pelz et al., 2017). Scale bars: 100 μm. (E) Vimentin (red) and DAPI (light blue) staining in testis paraffin sections of WT (α7^+/+^), importin α7^ΔIBB/ΔIBB^ and α7^−/−^ mice; scale bars: 100 μm. (F) H&E staining of testis paraffin sections showing aberrant sperm orientation in seminiferous tubules of importin α7^ΔIBB/ΔIBB^ and α7^−/−^ testes. Arrows mark the misorientated sperms. (G) DAPI and PNA staining in testis paraffin sections of WT (α7^+/+^), importin α7^ΔIBB/ΔIBB^ and α7^−/−^ mice showing defective sperm transport through the seminiferous epithelium in importin α7^ΔIBB/ΔIBB^ testes. Arrows mark the mislocalized sperms. (H) Sperm retention in stage IX seminiferous tubules. To label sperm cells, immunohistochemistry using anti-acetyl-Histone H4 (H4K8Ac, green), PNA (red) and DAPI (light blue) was performed. Asterisks indicate sperm retention. Scale bars: 25 μm.

Further analysis of Sertoli cell cytoskeletal proteins revealed an abnormal localization of the intermediate filament vimentin in Sertoli cells of importin α7^ΔIBB/ΔIBB^ and importin α7^−/−^ mice. Vimentin-based filaments no longer stretched across the Sertoli cell cytosol but retracted from the cell cortical region and were wrapped around the cell nuclei (Fig. 6E). In contrast, beta-III tubulin organization was not perturbed in Sertoli cells of importin α7^ΔIBB/ΔIBB^ and importin α7^−/−^ mice (data not shown), suggesting that there is no general effect on Sertoli cell morphology but rather a specific change in vimentin distribution.

The compromised Sertoli cells lead to defects in sperm orientation in both importin α7^ΔIBB/ΔIBB^ and importin α7^−/−^ mice. (Fig. 6F). Interestingly, sperm transport through the seminiferous epithelium, which is also dependent on Sertoli cells, was severely disturbed in importin α7^ΔIBB/ΔIBB^ mice, but was found to be unaffected in importin α7^−/−^ mice (Fig. 6G). In WT testis, only one generation of spermatids was found in stages IX-XII, during the transition from round into elongating spermatids. However, two generations of spermatids were often observed in stages IX-XII in importin α7^ΔIBB/ΔIBB^ mice (Fig. 6H). The additionally found spermatids were more mature with more condensed nuclei, which implicates that they were not released from Sertoli cells in the previous spermatogenesis cycle. Although correct spermiation is dependent on Sertoli cells, we could not detect residual sperms in stage IX-XII seminiferous tubules of importin α7^−/−^ testes, which excludes that spermatid persistence is caused by Sertoli cell defects only.

### Importin α7 deficiency-related loss of spermatocytes starts with leptotene/zygotene transition

To elucidate the start of germ cell loss during spermatogenesis in testes deficient for importin α7, we quantitatively evaluated different developmental steps of spermatogenesis. By labelling of spermatogonia with the pluripotency marker Sall4, no differences between genotypes were detected, excluding a severe loss of spermatogonia (Fig. 7A). In the adult testis the BrdU-positive cells in stages VII-VIII are preleptotene spermatocytes (Zhou et al., 2008). We observed that the number of BrdU-labeled preleptotene spermatocytes per tubule in importin α7^ΔIBB/ΔIBB^ and α7^−/−^ testes were similar to those in WT mice (Fig. 7A), indicating that importin α7 is dispensable for the development of preleptotene spermatocytes. We next tested for γH2AX, a phosphorylated form of histone 2AX, that exhibits an intense diffuse staining pattern in spermatocytes at the leptotene/zygotene transition in stages X-XI, and exclusively localizes to the sex chromosomes (so called sex body) within pachytene spermatocytes (Blanco-Rodriguez, 2009; Celeste et al., 2002; Peters et al., 1997). In adult importin α7^ΔIBB/ΔIBB^ testes and importin α7^−/−^ testes, the numbers of leptotene/zygotene spermatocytes in stages X-XI decreased markedly compared to WT controls while we did not observe differences in the stage-specific appearance of γH2AX-positive chromatin (Fig. 7A, B). Interestingly, we detected a further decrease in the number of stage I-VIII pachytene spermatocytes in importin α7^ΔIBB/ΔIBB^ but not in importin α7^−/−^ testes showing that importin α7^ΔIBB/ΔIBB^ testes were more affected than importin α7^−/−^ testes (Fig. 7A). The ratios of pachytene to leptotene spermatocytes were similar in WT and importin α7^−/−^ testes, while it was markedly reduced in importin α7^ΔIBB/ΔIBB^ mice (wt: 0.94; α7^ΔIBB/ΔIBB^: 0.77, α7^−/−^: 1.00), suggesting that development of pachytene spermatocytes is dependent on importin α7. The reduced numbers of step 1-8 round spermatids in importin α7^ΔIBB/ΔIBB^ and importin α7^−/−^ testes were comparable with numbers of pachytene spermatocytes (Fig. 7A), while the ratios of round spermatids to pachytene spermatocytes were similar between WT (2.6), α7^ΔIBB/ΔIBB^ (3.3) and α7^−/−^ testes (3.2). These observations suggest that deficiency of importin α7 leads to a reduction in leptotene/zygotene spermatocytes, and to a further decrease in pachytene spermatocytes, but surviving spermatocytes could differentiate into round spermatids.

**Fig. 7.**
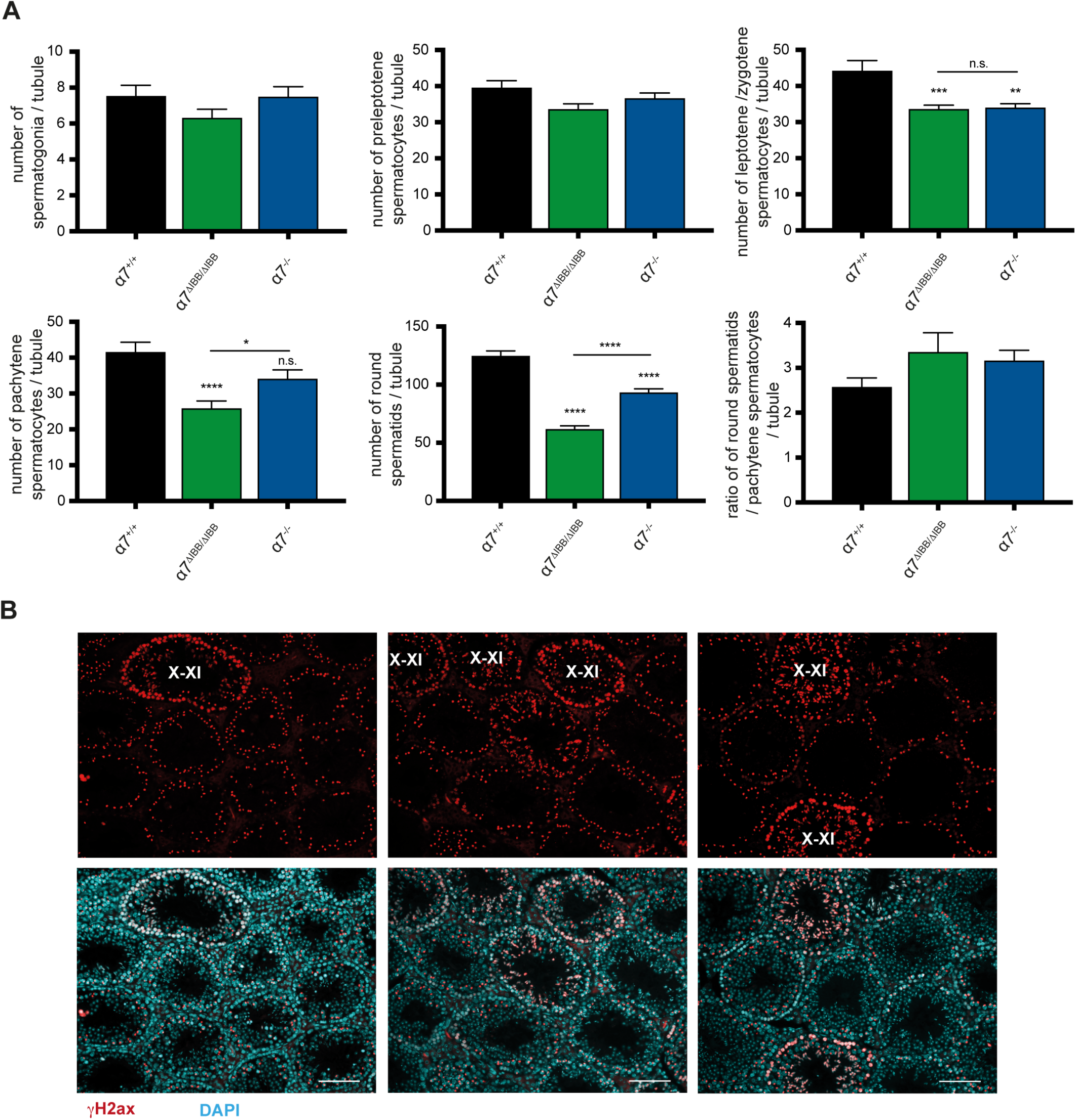
Importin α7 deficiency results in a loss of spermatocytes. (A) Number of spermatogonia, preleptotene, leptotene/zygotene, pachytene spermatocytes and round spermatids per seminiferous tubule in adult WT (α7^+/+^), importin α7^ΔIBB/ΔIBB^ and α7^−/−^ testes. (B) γH2AX (red) and DAPI (light blue) staining in adult WT, importin α7^ΔIBB/ΔIBB^ and α7^−/−^ testes. X-XI: stage X-XI tubuli, scale bars: 100 μm.

### Onset of spermatogenesis is delayed in importin α7^ΔIBB/ΔIBB^ mice

By H&E staining of testes from 6 weeks old mice we discovered a delayed onset of the first wave of spermatogenesis: while WT and importin α7^−/−^ mice showed a regular histology with seminiferous tubules at various stages, in importin α7^ΔIBB/ΔIBB^ mice, all tubuli displayed uniformly the same developmental stage and no round spermatids or later stages could be detected, indicating that the first meiotic wave had not been completed (Fig. 8A). We investigated this further with BrdU/γH2AX co-labeling in day 14 mouse testes. Since BrdU labels mitotic spermatogonia and preleptotene spermatocytes (Zhou et al., 2008) and, γH2AX stains B-type spermatogonia and preleptotene spermatocytes (Blanco-Rodriguez, 2009; Hamer et al., 2003), the double-stained germ cells should be B-type spermatogonia and preleptotene spermatocytes. We observed early pachytene spermatocytes (γH2AX confined to the sex chromosomes) in many tubules of WT testes, while very few tubules contained early pachytene spermatocytes in α7^ΔIBB/ΔIBB^ mice (Fig. 8B). In WT testes, we found only few double-stained preleptotene spermatocytes; in contrast many preleptotene spermatocytes showed up in the α7^ΔIBB/ΔIBB^ tubules. These observations suggested that in day 14 testes, most of the WT germ cells already had passed through the preleptotene spermatocyte stage, and many of them progressed into early pachytene spermatocyte stage, however most of the mutant germ cells remained at preleptotene spermatocyte stage, with only few of them reaching pachytene stage (Fig. 8B). Analysis of day 21 testes by γH2AX labelling revealed a high number of leptotene/zygotene spermatocytes in importin α7^ΔIBB/ΔIBB^ mice, while in WT testes most of the spermatocytes had already reached pachytene stage and round spermatids start to be present (Fig. 8C). Together, these data suggest that, compared with WT mice, the onset of the first wave of spermatogenesis is markedly delayed in α7^ΔIBB/ΔIBB^ mice.

**Fig. 8.**
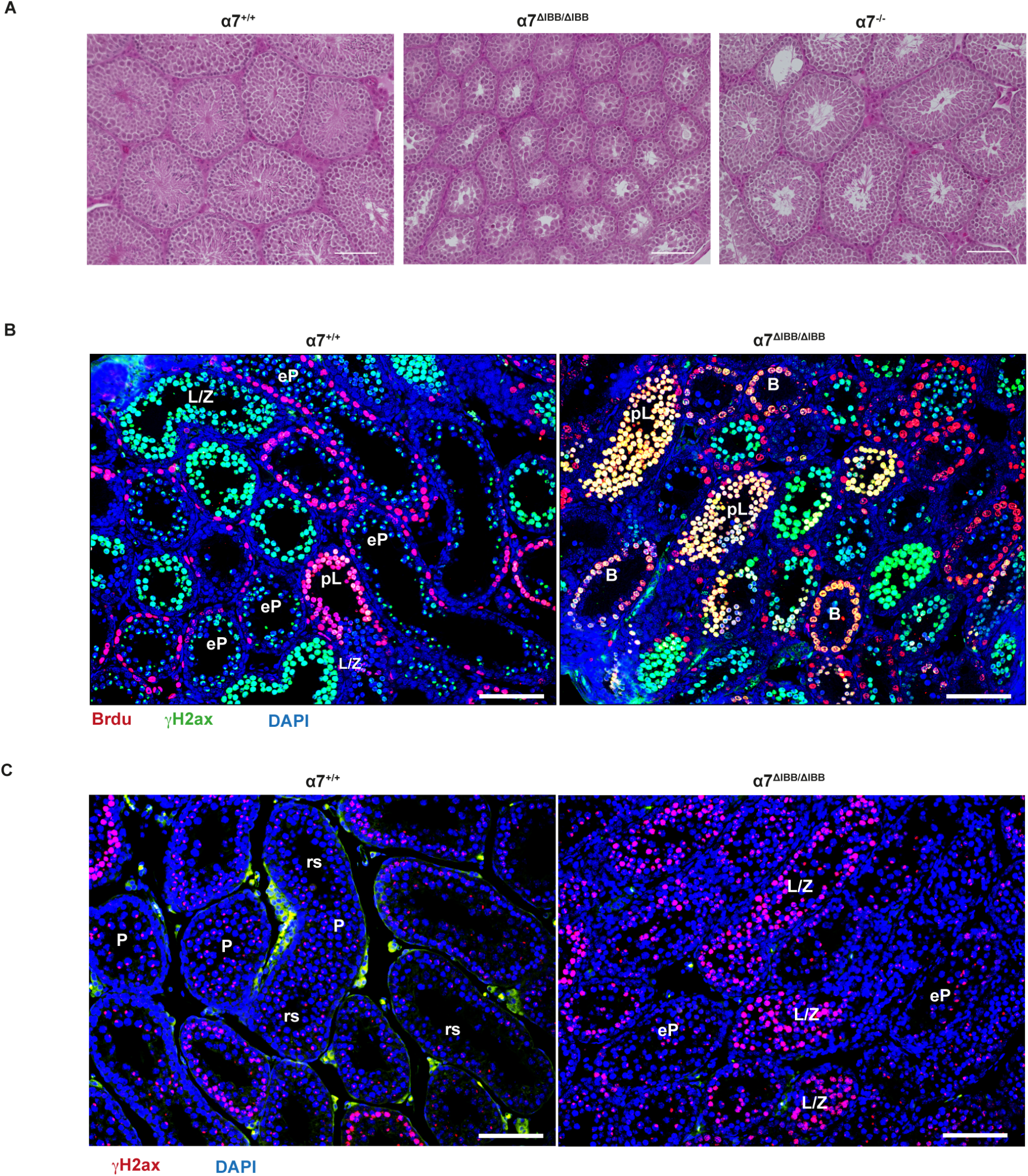
Importin α7 deficiency causes a delay in the first spermatogenesis wave. (A) H&E staining of testis paraffin sections of WT (α7^+/+^), importin α7^ΔIBB/ΔIBB^ and α7^−/−^ at 6 weeks of age showing severely delayed onset of spermatogenesis. (B) BrdU (red), γH2AX (green) and DAPI (blue) staining on paraffin sections of day 14 WT (α7^+/+^) and importin α7^ΔIBB/ΔIBB^ testes. The yellow colour indicates colocalization of BrdU and γH2AX (C) γH2AX (red) and DAPI (blue) staining on paraffin sections of day 21 WT and importin α7^ΔIBB/ΔIBB^ testes. B: B spermatogonia; pL: preleptotene spermatocytes; L/Z: spermatocytes at leptotene to zygotene transition; eP: early pachytene spermatocytes; P: pachytene spermatocytes; rs: round spermatids. Scale bars: 100 μm.

### Gene expression changes in importin α7^ΔIBB/ΔIBB^ testis

To obtain a transcriptome-wide insight into the affected transcripts, pathways and upstream regulators after Kpna6 depletion, we performed an RNAseq analysis on whole testes of WT, importin α7^ΔIBB/ΔIBB^ and α7^−/−^ mice in triplicate. A principal component analysis on the gene aggregated transcript per million values depicts the transcriptome of the importin α7^ΔIBB/ΔIBB^ relative to the WT and the importin α7^−/−^ mice (Fig. S2). According to the PCA the transcriptomes of the former differ the most from the latter two along the first principal component (PC1), which is in accordance with the much more severe phenotype observed in importin α7^ΔIBB/ΔIBB^ compared to importin α7^−/−^ mice. A log likelihood test between the importin α7^ΔIBB/ΔIBB^ and the WT transcriptome revealed 112 significantly regulated genes (p-value < 0.01; absolute effect size > 1), with Kpna6 significantly downregulated in the former (p-value: 3*10^−5^; effect size: −1.51) (Fig. 9A; Table S3). Contrary to this and in line with the mild phenotype and the PCA, we found only 20 differentially regulated genes between WT and importin α7^−/−^ mice (p-value < 0.01; absolute effect size > 1), with Kpna6 showing no significance in differential expression. Therefore, we concentrated our analyses on the comparison of importin α7^ΔIBB/ΔIBB^ and WT testes. We next performed a gene set analysis based on the gene effect size differences between the WT and importin α7^ΔIBB/ΔIBB^ condition using the biological processes from the Gene Ontology (GO). Upregulated GO terms in importin α7^ΔIBB/ΔIBB^ are related to cell migration, extracellular matrix and development processes, while cilia, flagellum and sperm related terms are downregulated due to the dysfunctional Kpna6 gene (Fig 9B). To obtain an insight into the upstream transcription factors driving the respective gene expression, we tested the mouse regulon data from the DoroTheA library against the WT and importin α7^ΔIBB/ΔIBB^ condition. In total there were 16 (20) transcription factors (TF) whose putative activity was significantly up- (down-) regulated in the importin α7^ΔIBB/ΔIBB^ mice (Fig. 9C). While the upregulated TF were related to TNF-alpha signaling via NF-kB (adj. p-value=0.0008) and TGF-beta signaling (adj. p-value=0.02), Rfx2, a key regulator of mouse spermiogenesis was downregulated. To further test the prediction, we compared the differential gene regulation from our importin α7^ΔIBB/ΔIBB^ mice with testicular transcriptomes of Rfx2 knockout mice (GEO ID GSE68283)(Wu et al., 2016). The effect size and the direction of differential gene regulation for Rfx2 knockout and α7^ΔIBB/ΔIBB^ mice relative to the wild type conditions are significantly correlated according to Spearman’s rho statistic (p-value < 2.2*10^−^ 16). While both mouse lines share few upregulated genes (10% of Rxf2 KO; 7% of Kpna6 KO), 40% of the genes that were downregulated in Kpna6 KO, were also downregulated in Rfx2 KO (Fig. 9D, Table S4). Among the commonly downregulated genes are various proteases of the ADAM family, which are known to be involved in sperm function (Cho, 2005). A GSEA of the downregulated pathways in Rfx2 knockout and α7^ΔIBB/ΔIBB^ mice depicted similar effects on cilium, its assembly, microtubule- and sperm motility-related processes (Fig.9E, Table S5). Pathways that are significantly downregulated in importin α7^ΔIBB/ΔIBB^ mice (p-value < 0.05), but not in Rfx2 KO mice (p-value > 0.1) include chemical synaptic transmission, pyroptosis, sperm principal piece and acrosomal vesicle and hydrolase activity.

**Fig. 9.**
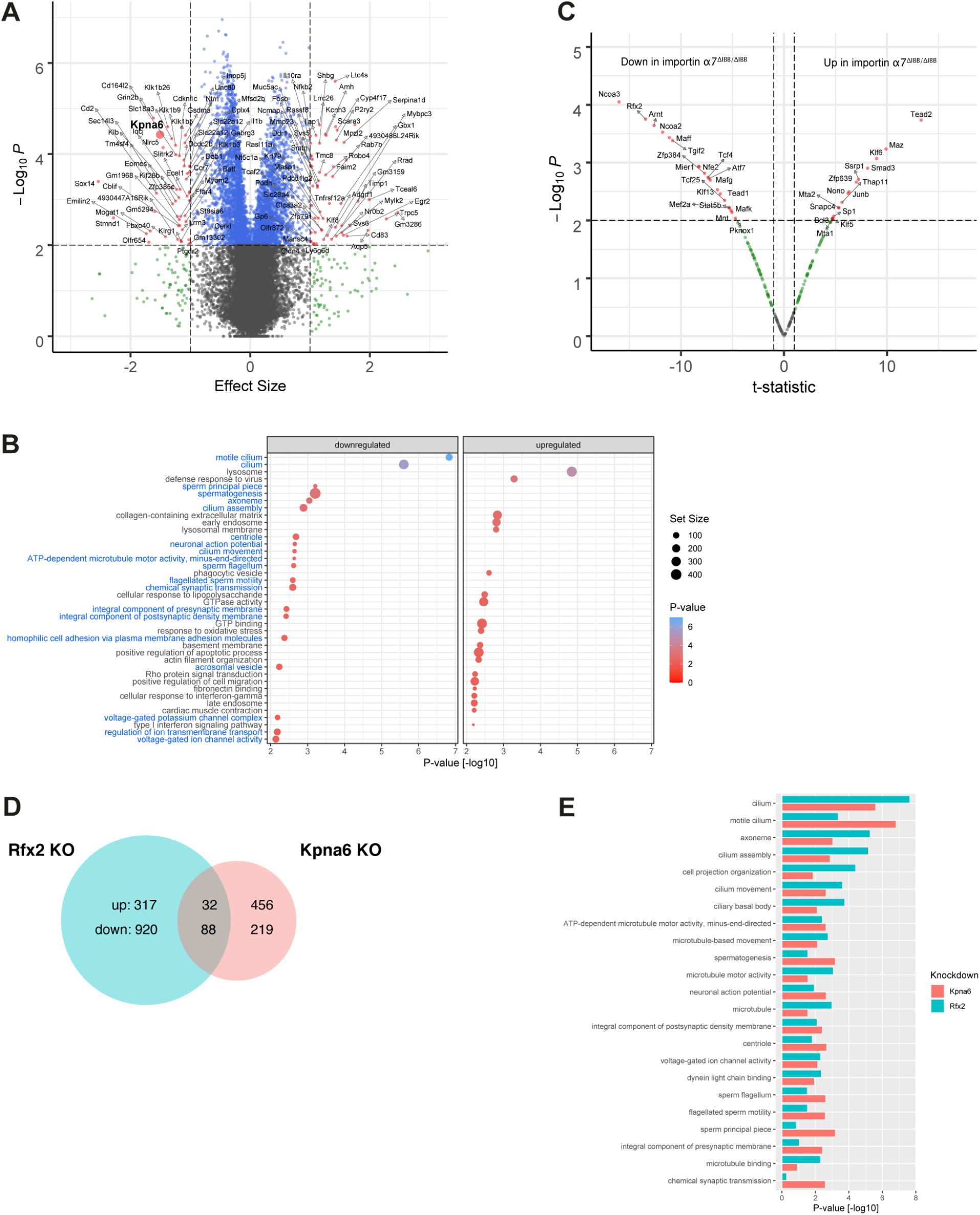
Downregulated spermiogenesis-related pathways in importin α7-deficient testis. (A) Volcano plot depicting the differential gene regulation of the testes transcriptomes of WT and importin α7^ΔIBB/ΔIBB^ mice. The x- and the y-axes show the effect size according to a Wald test and the negative log_10_ transformed p-value. Small red dots indicate significantly regulated genes according to the p-value (p< 0.01) and the absolute effect size (>1). The Kpna6 gene is indicated by an enlarged red circle. (B) Dotplot depicting the 20 most significantly up- and downregulated GO terms in the testes transcriptomes of importin α7^ΔIBB/ΔIBB^ mice compared to WT. The locus and color of the dots indicates the −log_10_ p-value, while the dot size is related to the number of genes of the GO term. Blue and black row names refer to the down- and upregulated GO terms, respectively. (C) Volcano plot depicting the predicted differential TF activity of the testis transcriptomes of WT and importin α7^ΔIBB/ΔIBB^ mice. The x- and the y-axes show the t-statistic of a t-test and the negative log10 transformed p-value. Red dots indicate significant differential TF activity according to the p-value (p< 0.01) and the absolute t-statistic. (D) Venn diagramm showing the commonly and individually up- and downregulated genes from mouse testes in a Rfx2 KO and the importin α7^ΔIBB/ΔIBB^ model in comparison to their respective wildtype controls. P-value cutoff < 0.01, absolute effect size > 0.5. (E) Barplot depicting simultaneously the significance of 23 most downregulated GO terms in testes transcriptomes after importin α7 depletion (Kpna6, red) or Rfx2 knockdown (GEO ID GSE68283) (Rfx2, green) according to a gene set enrichment analysis. The bars denote the −log_10_ transformed p-values.

### Downregulation of protamines and transition proteins in round spermatids and impaired histone-protamine exchange in importin α7^ΔIBB/ΔIBB^ mice

Since analysis of the altered gene expression in importin α7^ΔIBB/ΔIBB^ mice suggested the strongest impact on postmeiotic events, we performed a detailed microscopic study of sperm maturation in the testis. While spermatid elongation started regularly in stage IX seminiferous tubules of importin α7^ΔIBB/ΔIBB^ mice, the subsequent steps were characterized by abnormal nuclear condensation of elongating spermatids in importin α7^ΔIBB/ΔIBB^ mice compared to WT and α7^−/−^ testes (Fig. 10) and mature step 15-16 sperms were absent in importin α7^ΔIBB/ΔIBB^ mice. Together with the reduced number of spermatozoa and abnormal morphology of the epididymal sperms in α7^ΔIBB/ΔIBB^ mice this suggested that spermiogenesis was severely affected by importin α7 deficiency, which confirmed the results of gene expression analysis. We next tested whether the regulation of post-meiotic factors, that are necessary for sperm maturation, is mediated by importin α7. The analysis of gene expression in whole testis had already shown a slight downregulation of Tnp1, Tnp2, Prm1 and Prm2 (respective adj. p-values: 0.007, 0.06, 0.06, 0.02; Table S3) Real-time PCR analysis now demonstrated that the mRNAs of *tnp1*, *tnp2*, *prm1* and *prm2* were significantly reduced in isolated round spermatids of importin α7^ΔIBB/ΔIBB^ mice compared to WT (Fig. 11A). Thus, the reduced detection of protamines and transition proteins in importin α7^ΔIBB/ΔIBB^ is partly due to decreased expression of these genes in round spermatids. In contrast, other postmeiotic key genes were not significantly affected in importin α7^ΔIBB/ΔIBB^ mice (Fig. 11A). Moreover, protein analysis of chromatin-bound proteins showed that Tnp2 was drastically decreased in the testes of α7^ΔIBB/ΔIBB^ mice (Fig. 11B). Yet, Tnp1 and Tnp2 immunofluorescence revealed a regular localization of these two proteins in nuclei of elongating spermatids starting at step 9 (data not shown). However, while in WT Tnp1 cannot be found in tubules later than stage I, and Tnp2 is not expressed past stage III, we constantly found Tnp1-positive sperms until stage III tubules und expression of Tnp2 was even found in sperms of stage VIII (and IX, residual sperms) tubules, suggesting a severe disturbance of protamine-histone exchange in these sperms (Fig.11C, D).

**Fig. 10.**
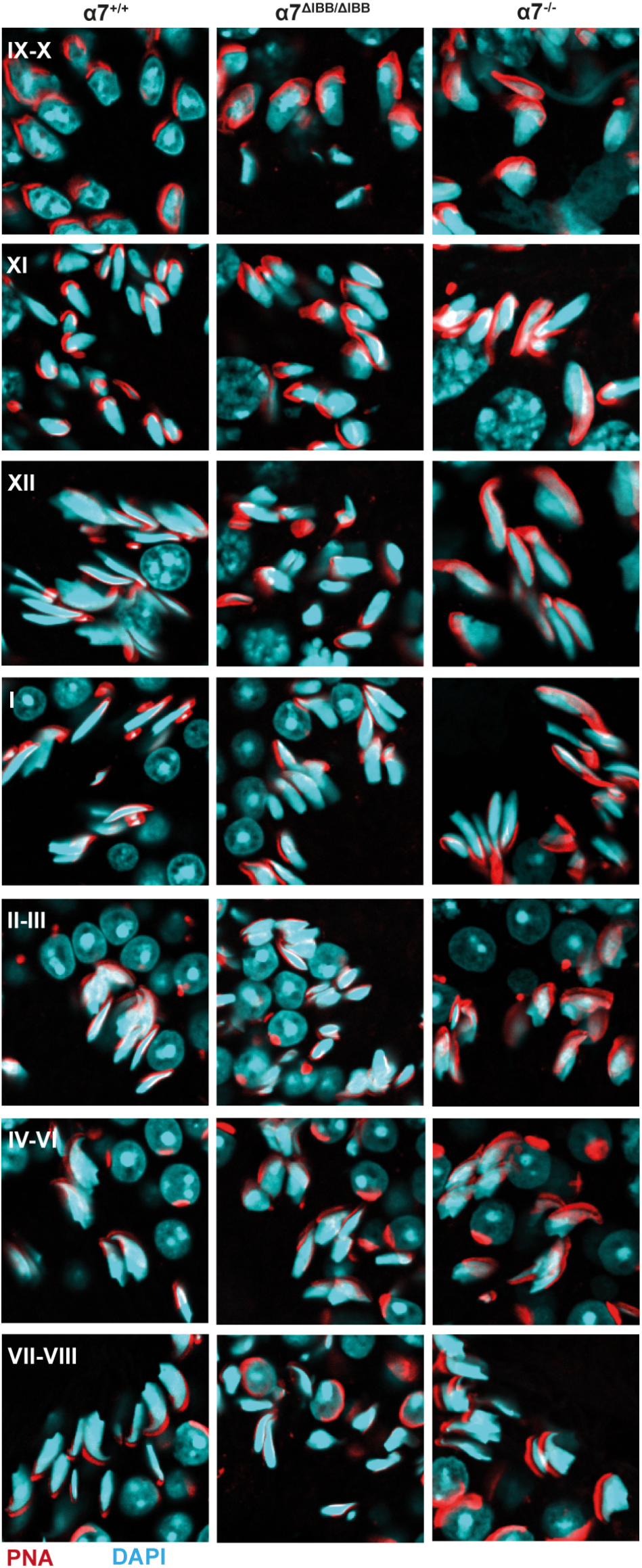
Importin α7-deficient spermatids show abnormal elongation and nuclear condensation. Morphology of DAPI- (blue) and PNA- (red) labelled spermatids throughout their development starting with elongation (step9). The deficiency/delay in elongation can be seen already in stage XII tubuli. Note the abnormal configuration of sperm heads in importin α7^ΔIBB/ΔIBB^ spermatids.

**Fig. 11.**
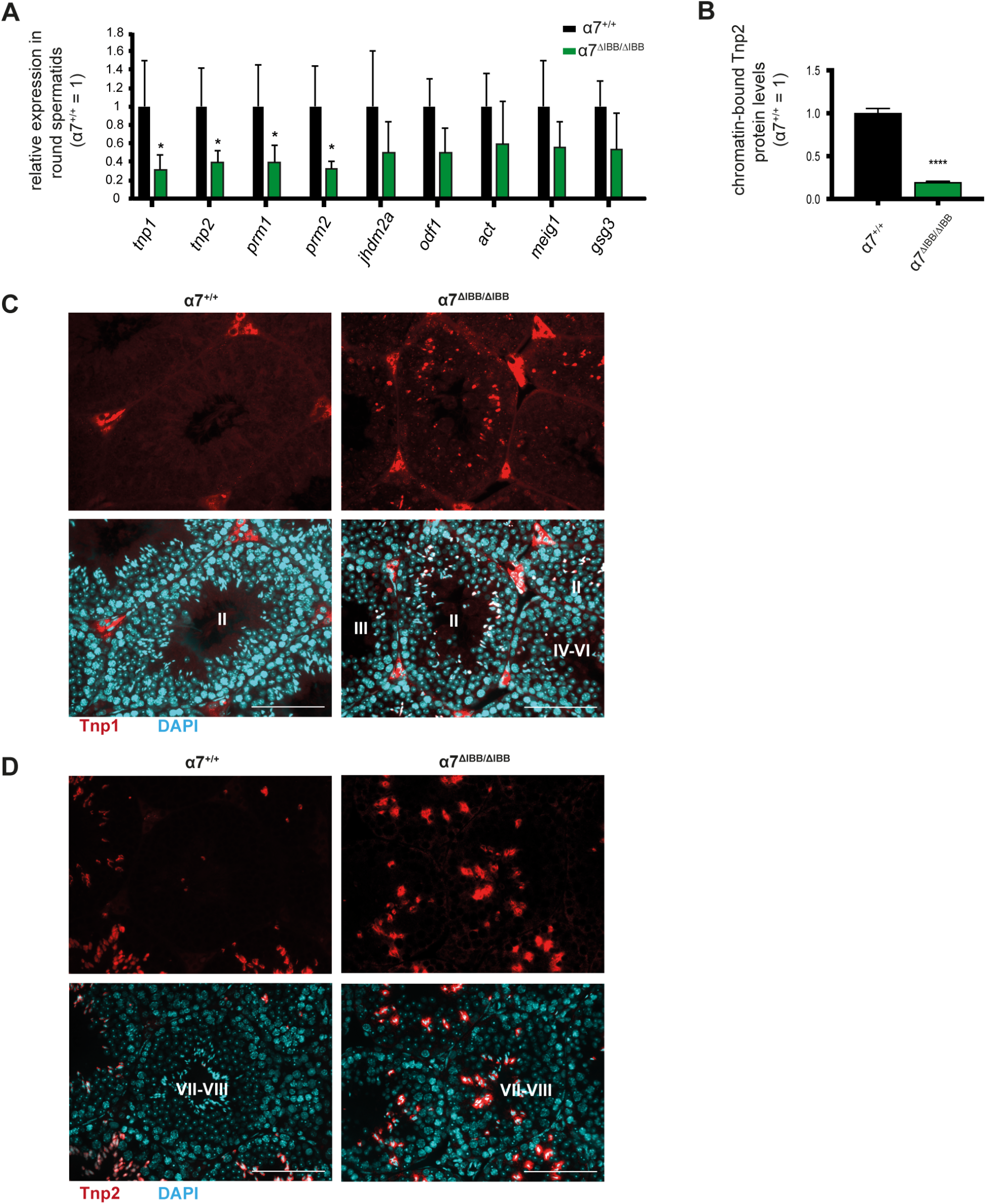
Reduced expression of transition nuclear proteins and protamines in importin α7-deficient testes. (A) Quantitative realtime PCR analysis of the expression levels of various postmeiotic genes including protamines and transition nuclear proteins in isolated round spermatids of adult WT (α7^+/+^) and importin α7^ΔIBB/ΔIBB^ testes (n=3-9). (B) Western blot analysis of Tnp2 in WT (α7^+/+^) and importin α7^ΔIBB/ΔIBB^ testis protein extracts. (C, D) Tnp1 (C, red), Tnp2 (D, red) and DAPI (light blue) staining on testis paraffin sections of adult WT and importin α7^ΔIBB/ΔIBB^ testes. Roman numbers mark the stage of seminiferous tubules; scale bars: 100 μm.

### Analysis of chromatin remodeling and presence of transcription factors in importin α7^ΔIBB/ΔIBB^ mice

Next we investigated the tremendous remodeling of chromatin that takes place during spermatid development, as this process is dependent on proteins that have to enter the nucleus of round and elongating spermatids. Analysis of histone H3 2- and 3-methylation as well as hyperacetylation of histone H3 (lysin 9 and lysin 14) and H4 (lysin 8 and lysin 12) did not reveal any differences of importin α7^ΔIBB/ΔIBB^ and WT spermatids (data not shown). Moreover, we observed regular appearance of DNA double strand breaks as detected by γH2AX labelling (Fig. 7B).

The significant reduction in gene expression of *tnp1, tnp2, prm1* and *prm2* may result from impaired expression or nuclear translocation of transcription factors. In mice, the transcriptional activation of these genes is mainly regulated by the transcription factor cAMP-response element modulator (Crem) (Mali et al., 1989; Nantel et al., 1996). WT testis showed a normal expression and localization of Crem in round spermatids from stage II-VIII (Fig. S3). With the onset of elongation the Crem expression started to decline. However, no major changes in Crem expression and localization could be observed in importin α7^ΔIBB/ΔIBB^ testis (Fig. S3), suggesting that nuclear import of Crem is not affected.

The transcriptional regulator Brwd1 has recently been shown to be essential for spermiogenesis (Pattabiraman et al., 2015). Being part of a postmeiotic transcriptional activator complex, Brwd1 binds to acetylated lysine residues of histones, causing transcriptional activation. Immunofluorescence of Brwd1 revealed striking differences in its expression in importin α7^ΔIBB/ΔIBB^ testes compared to WT testes. While in WT, positive signals could be found within paraffin embedded testis sections in step 9 as well as in step 14-15 elongating spermatids, no Brwd1 was found in importin α7^ΔIBB/ΔIBB^ testis. As the signals tended to be very subtle in paraffin sections of the testis, immunostaining was repeated in snap-frozen sections, showing very clearly the presence of Brwd1 in WT, but not in importin α7^ΔIBB/ΔIBB^ testes (Fig.12 A, B).

**Fig. 12.**
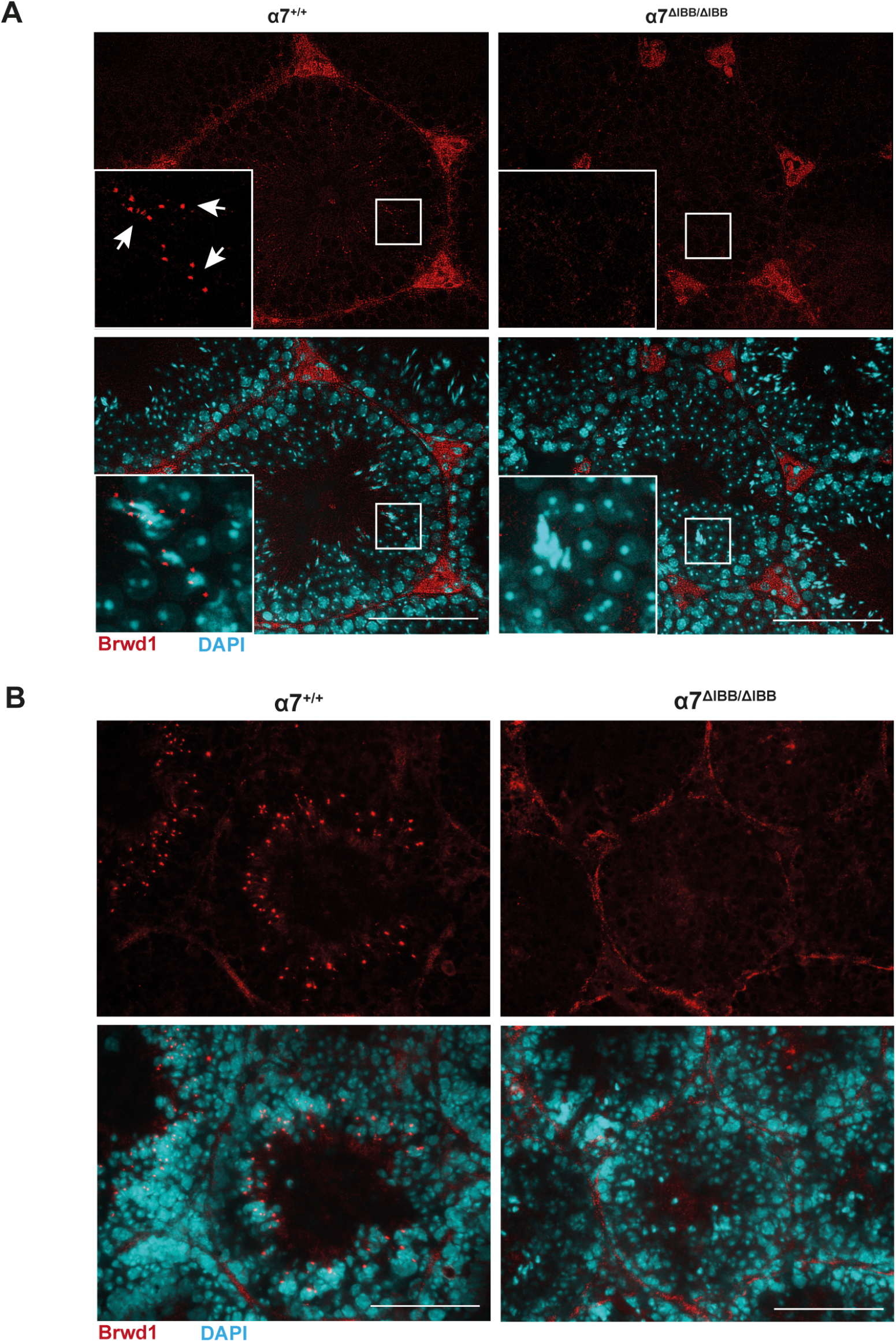
Disturbed expression of transcriptional regulator Brwd1 in importin α7-deficient testes. (A, B) Immunofluorescence for Brwd1 (red) in importin α7^+/+^ and α7^ΔIBB/ΔIBB^ testis on paraffin sections (A) or on snap-frozen cryosections (B); counterstain with DAPI (light blue), scale bars: 100 μm.

## Discussion

In the present study, we analyzed the biological function of importin α7 (KPNA6) during spermatogenesis using two different knockout mouse models, and we showed for the first time that importin α7 is essential for mammalian male fertility. These findings are in accordance with the phenotype of fruit flies lacking importin α1, which is the one of the three known importin α paralogs in *D. melanogaster* with the highest similarity to mouse importin α7. The importin α1 of *D. melanogaster* is essential for male gametogenesis exhibiting full arrest in spermatogenesis and complete male infertility (Ratan et al., 2008). The sterility effect was paralog-specific and resulted in defective spermatocytes with abnormal nuclear shape suggesting a disturbed chromatin condensation in these cells. However, the molecular mechanisms by which *D. melanogaster* importin α1 exerts its effects on male fertility have not been assessed yet.

Expression of importin α7 in mouse testis has been investigated at the mRNA level by several techniques. It has been shown by microarray that importin α7 mRNA expression is increased in isolated pachytene spermatocytes and round spermatids (Holt et al., 2007; Major et al., 2011; Shima et al., 2004). Importin α7 mRNA is upregulated during the first spermatogenesis wave: a first increase takes place from postnatal day 10 to day 20, when spermatocytes progress into late pachytene stage; and a second increase occurs from postnatal day 20 to day 30, when the round spermatids become enriched (Major et al., 2011; Namekawa et al., 2006). The importin α7 protein expression pattern in germ cells during spermatogenesis detected by immunohistochemistry in our current report is consistent with these data. In addition, microarray data had shown that importin α7 mRNA is moderately expressed in *in vitro* cultured Sertoli cells isolated from postnatal day 16 to day 18 testes (Holt et al., 2007; Shima et al., 2004), while in adult testis, importin α7 mRNA was not detected in Sertoli cells by *in situ* hybridization (Hogarth et al., 2006). However, our present study shows, that in addition to germ cells, importin α7 protein is also highly expressed in the nuclei of Sertoli cells throughout all stages of the seminiferous epithelium in the adult testis.

Studies of other importin α isoforms in adult mouse testis have shown stage- and cell type-specific expression of importin α1 α3, α4 and α5 (Miyamoto et al., 2012). The distinct expression patterns of importin α proteins in testis imply that they may play defined roles during mammalian spermatogenesis We confirmed the published expression of α-importins in murine testis, and using specific antibodies for each importin α paralogue, our own comprehensive studies revealed, that importin α7 is the only importin α isoform expressed in elongating spermatids. Moreover, we detected a massive increase in importin α7 expression in late round and early elongating spermatids that has not been seen for any other importin α subtype. The infertility phenotype of importin α7-deficient mice underlines for the first time the hypothesis that this protein has unique functions in spermatogenesis, and other importin α subtypes cannot compensate for its absence.

One of the main findings of our present work is that importin α7 has two different functions during spermatogenesis depending on its two localizations in Sertoli cells and in developing sperms. The use of two different importin α7-deficient mouse lines with different expression patterns in these cell types allowed us to discriminate between these two functions as the two mouse lines display distinct phenotypes. The importin α7^ΔIBB/ΔIBB^ mice are infertile and are characterized by the absence of the full-length protein in Sertoli cells as well as in developing sperms. On the other hand, importin α7^−/−^ mice are fertile and produce sperms and this can be clearly attributed to a rescued expression of the protein in developing sperms based on a testis specific promoter in intron 1 of the *kpna6* gene. The utilization of a testis-specific promoter is a well-known mechanism in male germ cells and has been shown for a variety of transcripts encoding for proteins such as angiotensin converting enzyme, c-abl, proopiomelanocortin, superoxide dismutase or ß-galactosyltransferase (Hecht, 1998). In importin α7^−/−^ mice, germ cells, but not somatic cells of the testis produce importin α7 mRNA by using an alternative exon 1A, leading to mRNA with a germ cell specific 5’-untranslated region, suggesting differing transcriptional control mechanisms for importin α7 in germ cells and somatic cells. However, the absence of importin α7 in Sertoli cells seems to account for a partially reduced sperm count. This finding could be confirmed by intercrossing of both lines, producing compound heterozygous offspring, which also displayed a reduced sperm count but were fertile. The reduced sperm count in these heterozygous compound mice provides clear evidence that the partial sperm count reduction can be attributed to the Sertoli cell related phenotype, as the compound heterozygous mice represent a rescue of the infertile importin α7^ΔIBB/ΔIBB^ mice with expression of the full-length protein in germ cells, but not in Sertoli cells.

The functions of Sertoli cells include providing structural support and nutrition for developing germ cells, phagocytosis of degenerating germ cells and residual bodies, establishment of the BTB and release of spermatids at spermiation. In this study, we have shown that importin α7 is highly expressed in nuclei of Sertoli cells in WT testes. Additionally, we found the number of Sertoli cells to be reduced in the seminiferous tubules of importin α7^−/−^ and α7^ΔIBB/ΔIBB^ testes due to degeneration and shedding.

We observed a nuclear import deficiency of the AR in Sertoli cells of both mutant mouse lines. It has been shown that ligand-dependent nuclear import is crucial for the function of the AR in both health and disease (Becker et al., 2000; Kawate et al., 2005; Nakauchi et al., 2007; Thomas et al., 2004). The unliganded AR is retained in the cytoplasm but, on binding of 5α-dihydrotestosterone, it translocates into the nucleus and alters transcription of its target genes. The nuclear import of AR has been shown to be importin α/ß dependent (Cutress et al., 2008; Kaku et al., 2008). Mice with Sertoli cell-specific ablation of the AR exhibit defective Sertoli cell maturation and polarization, increased permeability of the BTB and downregulation of Cldn3, suggesting that Cldn3 is a transcriptional target of AR (Meng et al., 2011; Meng et al., 2005; Willems et al., 2010). We found several AR-dependent genes with the tendency towards an upregulation in importin α7^ΔIBB/ΔIBB^ mice and with a normal expression in importin α7^−/−^ mice. The apparent upregulation is predictable, as the loss of spermatids is much higher in α7^ΔIBB/ΔIBB^ mice than the loss of Sertoli cells, leading to a higher relative amount of Sertoli cells compared with WT testis.

Nevertheless, c*ldn3* mRNA levels were significantly decreased in importin α7^ΔIBB/ΔIBB^ testes while being normal in importin α7^−/−^ testes, suggesting that *cldn3* is regulated by additional mechanisms that differ between the two mutant lines; moreover, Chihara et al. found *cldn3* not only to be expressed by Sertoli cells, but mainly by Stra8 positive germ cells (Chihara et al., 2013), which could explain the differences found in importin α7^ΔIBB/ΔIBB^ and α7^−/−^ testes. The reduced expression of *cldn3* mRNA, however, does not affect the correct localization of the protein in stage VIII seminiferous tubuli in importin α7^ΔIBB/ΔIBB^ mice, although the quantitative amount of a protein is difficult to estimate by immunohistochemistry. It has been shown that during stages VII and VIII the BTB is dynamically remodeled as preleptotene spermatocytes pass through the barrier (Bremner et al., 1994; Dym and Fawcett, 1970). We observed a relative reduction in leptotene/zygotene spermatocytes of about 25% in importin α7^ΔIBB/ΔIBB^ and α7^−/−^ seminiferous tubules which would be consistent with a meiotic delay due to a prolonged leptotene/zygotene phase as it had been observed in *cldn3* knockdown mice (Chihara et al.). The presence of testis-specific autoantibodies in the blood of both mutant lines also suggests a disturbance of the BTB integrity which depends on Sertoli cells. However, we could not detect higher IgG levels in testes of both mouse lines compared to WT and *in vitro* biotin diffusion assays did also not confirm a permanently disturbed BTB. The contradictory results regarding BTB stability lead us to speculate that there is a partially compromised BTB, resulting in presence of autoantibodies and delayed passage of leptotene/zygotene spermatocytes. Although both mutant mouse lines exhibit a loss of leptotene/zygotene spermatocytes, only importin α7^ΔIBB/ΔIBB^ mice show an even more pronounced loss of pachytene spermatocytes and round spermatids, suggesting that the latter can be attributed to the defective importin α7 in germ cells. Although we detected defective sperm orientation in both mutant lines, which can be attributed to defective Sertoli cell function, the sperm transport through the epithelium was found to be dysregulated only in importin α7^ΔIBB/ΔIBB^ mice but normal in importin α7^−/−^ mice. The fact, that in α7^ΔIBB/ΔIBB^ testes stage IX-XII tubules frequently contain spermatids at different maturation steps leads to the conclusion that spermatids are not properly released from the seminiferous epithelium. We were not able to detect such residual spermatids in importin α7^−/−^ testis, suggesting that a failure in coordinated spermiogenesis rather than in Sertoli cell-dependent release accounts for the residual spermatids.

A striking finding of our present study was, that the Sertoli cells of both mutant lines display abnormal organization of the intermediate filament vimentin. While vimentin is stretched across the Sertoli cell cytosol in WT testis, both mutant mouse lines showed a concentration around the Sertoli cell nucleus with no extensions. Similar findings have been found in mice with a defective cytoplasmic dynein 1 heavy chain and in mice that have been depleted of raptor, a key component of mTORC1 (Wen et al., 2018; Xiong et al., 2018). However, in both of these mouse lines, the phenotype included a severely disorganization of actin and microtubules. We could not find a disordered localization of Sertoli cell microtubuli in importin α7^ΔIBB/ΔIBB^ or importin α7^−/−^ mice, excluding major morphological changes of Sertoli cell cytoplasm which still stretched out into the tubular lumen.

By H&E staining of 6 weeks old testes we found retardation in the first wave of spermatogenesis which was confirmed by detailed analyses in 14 and 21 days old testes. While early pachytene spermatocytes could be detected in day 14 WT testes, they were absent in importin α7^ΔIBB/ΔIBB^ mice. Moreover, at day 21, many tubules containing leptotene/zygotene spermatocytes were found in importin α7^ΔIBB/ΔIBB^ mice, while in WT testes most of the spermatocytes had already reached pachytene stage. The reason for this delay is not completely clear but it fits to previous reports of importin α7 being upregulated starting at postnatal day 10, suggesting a very specific role at this time point (Major et al., 2011; Namekawa et al., 2006).

Multiple molecular events have to occur for a round spermatid to become a mature sperm. These events include chromatin condensation, reorganization of the spermatid nucleus, formation of an acrosome and assembly of a sperm tail (Sassone-Corsi, 2002). It has been shown that a number of postmeiotic proteins including transition nuclear proteins 1 and 2 (Tnp1 and Tnp2), protamines 1 and 2 (Prm1 and Prm2), meiosis expressed gene 1 protein (Meig1), outer dense fiber protein 1 (Odf1), lysine-specific demethylase 3a (Jhdm2a), activator of Crem in testis (Act), and F-actin-capping protein subunit alpha-3 (Capza3), are important for spermiogenesis (Geyer et al., 2009; Kotaja et al., 2004; Liu et al., 2010; Okada et al., 2007; Salzberg et al., 2010; Yang et al., 2012; Zhang et al., 2009). The transition from round spermatids to mature spermatozoa was severely affected in α7^ΔIBB/ΔIBB^ mice, since the few spermatozoa that did form in α7^ΔIBB/ΔIBB^ animals (3% of WT levels) displayed abnormal heads. Spermiogenesis requires extensive chromatin condensation. Histone displacement, a process in which histones are initially replaced by Tnp1 and Tnp2 and subsequently by Prm1 and Prm 2, is required for chromatin condensation. Accordingly, the genetic ablation of transition proteins or protamines causes defective spermiogenesis (Cho et al., 2001; Yu et al., 2000; Zhao et al., 2004; Zhao et al., 2001) comparable to the phenotype of importin α7^ΔIBB/ΔIBB^ males. By RNAseq we found that importin α7 deficiency reduced *tnp1*, *tnp2*, *prm1*, and *prm2* mRNA expression in the testis and this effect was even more pronounced when isolated round spermatids were analyzed by quantitative real-time PCR. Moreover, total chromatin-bound Tnp2 was markedly reduced in testes of importin α7^ΔIBB/ΔIBB^ mice as assessed by western blot. Additionally, a prolonged localization of Tnp1 and Tnp2 was detected in spermatids of importin α7^ΔIBB/ΔIBB^ mice, confirming the defective histone/protamine exchange.

The transcriptional activation of Tnp and Prm genes is mainly regulated by the transcription factor Crem and transcriptional activity of Crem is regulated by Act (Fimia et al., 1999; Mali et al., 1989; Nantel et al., 1996). It has been demonstrated that the Crem-regulated gene expression pathway is essential for normal spermatogenesis (Deng and Lin, 2002; Giorgini et al., 2002; Kotaja et al., 2004; Nantel et al., 1996). Like importin α7, Crem and Act are also highly expressed in round spermatids (De Cesare et al., 2003; Delmas et al., 1993). Analysis of Crem expression by immunofluorescence, however, did not reveal a significant reduction or mislocalization in importin α7^ΔIBB/ΔIBB^ testis. Moreover, other above mentioned postmeiotic (Crem-regulated) genes did not show a significantly reduced expression in importin α7^ΔIBB/ΔIBB^ testes.

The transcriptional regulator Brwd1 has recently been shown to be essential for spermiogenesis (Pattabiraman et al., 2015). Although a distinct pathway has not been found yet, the altered transcription of postmeiotic genes in Brwd1-defective testes has been suggested to cause male infertility. We found a striking reduction of Brwd1 testicular signals in importin α7^ΔIBB/ΔIBB^ testes that could be causative for the infertility in this mouse line.

A gene set enrichment analysis of the RNAseq data clearly indicated a downregulation of cilium- and sperm motility related processes in importin α7^ΔIBB/ΔIBB^ mice, corroborating the finding of defective spermatogenesis. Upregulated pathways include lysosomal and early endosome GO terms, which might be a consequence of the decreased importin α7 function. An upstream analysis of putative transcription factors singled out a downregulation of Rfx2 (Regulatory Factor X2) gene targets. While the transcription factor itself was not differentially regulated relative to WT, its targets related to cilium, axoneme, microtubule and sperm motility were, which was evident from a comparison of our RNAseq data to a testis transcriptome of Rfx2 knockout mice (Wu et al., 2016). Thus, it might be possible that the translocation of Rfx2 into the nucleus could be affected in importin α7^ΔIBB/ΔIBB^ testes.

The well-known function of importin α7 as part of the nuclear import system raises the possibility that import deficiency of one or more factors contribute to the phenotype directing the attention towards chromatin modifying proteins and transcriptional regulators. Having investigated transcriptional regulators as CREM and Brwd1 during the course of this study, there are many more possible factors that could be potentially involved in the phenotype of importin α7-deficient mice. The TBP-related factor Trf2, which is abundantly expressed in testis, is essential for mouse spermiogenesis, just as well as Taf7l, that has been shown to cooperate with Trf2 to regulate expression of postmeiotic genes (Zhang et al., 2001; Zhou et al., 2013). In eukaryotic cells, the transcription of genes is carried out by RNA polymerase II. Depending on its need for transcription, the RNA polymerase II shuttles between nucleus and cytoplasm and this shuttling has been suggested to represent a mechanism of transcriptional control in particular in spermatogenesis (Custodio et al., 2006).

A large number of studies have identified spermiogenesis-relevant genes that function in posttranscriptional control in concert with importin α. In the developing male gamete RNA synthesis terminates during mid-spermiogenesis, which is long before the spermatid completes its differentiation into the spermatozoon, making a coordinated translational regulation of stored mRNAs essential, and the formation of respective RNA-protein complexes has previously been shown to be regulated by importin α (Hecht, 1998; Sato and Maquat, 2009). The importance of such RNA-regulating proteins for a well-ordered germ cell development is underlined by an arrest at early spermiogenesis in mice lacking Piwi which regulates germ cell expressed miRNAs and piRNAs involved in mRNA silencing (Deng and Lin, 2002; Grivna et al., 2006). A further example is MSY2, a DNA/RNA binding protein solely expressed in germ cells, which has been implicated in stabilization of transcripts and translational repression. MSY2-deficient mice show a spermatogenic arrest in late round spermatids and infertility like importin α7^ΔIBB/ΔIBB^ mice (Yang et al., 2007).

The cap binding proteins CBP20 and CBP80 bind in the nucleus to newly formed transcripts and regulate their splicing, 3’-end formation, and nuclear export as well as the pioneer round of translation. The binding of CBP20 and CBP80 to mRNA, in turn, is tightly regulated by importin α and ß providing a further mechanism how importins could be involved in posttranscriptional control in spermatogenesis (Dias et al., 2009). It will be interesting to execute detailed analyses of these mechanisms in importin α7-deficient mice.

Taken together, in the present study we have shown for the first time that importin α7 is essential for mammalian male fertility. Importin α7-deficient testes exhibit a disturbed spermatogenesis with incomplete transition from preleptotene to leptotene spermatocytes and defective progression of round spermatids to mature spermatozoa. Using two different importin α7-deficient mouse lines for our analyses, we could clearly relate the above described phenomena to the missing expression of importin α7 in Sertoli cells and elongating spermatids, respectively. The compromised function of Sertoli cells is correlated with a partial decrease of germ cells and results in reduced sperm count. In contrast, impaired expression of importin α7 in round and elongating spermatids leads to complete arrest of spermiogenesis resulting in severe oligozoospermia. Since at the level of round spermatids importin α7 is the main expressed α-importin, its deficiency has rather pleiotropic effects, including transcriptional processes, proper localization of proteins and development of the sperm flagellum and shows its biggest impact during spermiogenesis when spermatids start to elongate. Although we revealed severe impairments in expression and localization of transition nuclear proteins and protamines, as well as of the transcription factor Brwd1, and altered expression of Rfx2 target genes, further investigations have to be performed to identify all the cellular and molecular mechanisms involved in the complex phenotype of infertility observed in α7^ΔIBB/ΔIBB^ mice.

## Materials and Methods

### Animals

Importin α7^−/−^ and α7^ΔIBB/ΔIBB^ mice were generated as described previously (Rother et al., 2011). Animals were backcrossed for 10 generations to C57Bl6/N background. If not stated otherwise, male mice aged 10-20 weeks were used for the analyses. All experiments were performed according to national and institutional guidelines and were approved by the relevant authority [Landesamt für Gesundheit und Soziales (LaGeSo), Berlin, Germany].

### Western blot

Mouse testes were collected and homogenized in RIPA-Buffer supplemented with protease inhibitor cocktail (Sigma-Aldrich, St. Louis, USA). Following sonication and centrifugation, protein concentrations in the tissue extracts were measured by using Bicinchorinic Acid Solution / Coppersulfate Solution 50:1 (Sigma-Aldrich, St. Louis, USA). 40 μg of total protein was separated by SDS-PAGE. After the transfer of proteins, the PVDF membrane was blocked by Odyssey blocking solution (LiCor, Bad Homburg, Germany) and subsequently incubated with primary antibodies at 4°C overnight. On the next day, the membrane was incubated with an IRDye-coupled secondary antibody for 1 h at room temperature and detection was performed using the Odyssey Infrared Scanner (LiCor, Bad Homburg, Germany). Signals were quantified using the Odyssey Infrared Scanner software (LiCor, Bad Homburg, Germany). The generation of C-terminal and N-terminal antibodies against importin α7 was accomplished using standard protocols and has been described previously (Kohler et al., 1999). The complete list of antibodies and conditions is found in supplemental data (Table S1).

### RNA-isolation, reverse transcription and PCR

Murine liver and testis were homogenized in trizol (FIRMA) and extracted with chloroform, subsequently precipitated in isopropanol and washed with 70% ethanol. The pellet was dried and resuspended in DEPC water. Then RNA was digested with DNAse I, and 2 μg of digested RNA was subjected to reverse transcription using a standard protocol. PCR was performed using 10 ng of cDNA, the used primers are listed in supplemental data (Table S2).

### Testosterone measurement

Blood samples were collected from 10-weeks-old mice by cardiac puncture. Serum samples were prepared as described earlier (Jeyaraj et al., 2005). The concentrations of testosterone in the serum samples were measured by using a Testosterone EIA kit (Cayman Chemical Company, Michigan, USA).

### Histological analysis

Testis and Epididymis were fixed in neutral buffered 4% formalin. After fixation, tissues were dehydrated in increasing concentrations of ethanol, embedded in paraffin wax, and sectioned at a thickness of 5 μm. Sections were deparaffinised, rehydrated and stained with hematoxylin and eosin according to standard protocols. For quantification of diameters of seminiferous tubules, images of H&E stained testis sections (5 animals per group) were taken using Keyence microscope (Keyence, Bioreva BZ-9000, Germany) and analyzed; 30 tubules per animal were measured using ImageJ software.

### Epididymal sperm count

Sperm count was performed as described previously (Liu et al., 2010; Wu et al., 2000). Briefly, one caudal epididymis was used for histological examination, and the other was minced in 1 ml of PBS. Sperms were allowed to disperse into solution by incubating for 5 min at room temperature. An aliquot of the sperm/saline mixture was then counted in a hemocytometer. The hemocytometer count was multiplied by appropriate volume and dilution factors to give a total cauda epididymal sperm count. The average number of sperms per epididymis was calculated from 4-8 mice for each genotype group.

### Immunohistochemical analysis

For IF staining sections underwent deparaffination followed by rehydration and antigen retrieval using either citrate buffer pH6 or Tris-EDTA buffer pH9 for 20 minutes, wherever appropriate. The sections were then treated with 10% normal donkey serum for 1h at room temperature and subsequently incubated with primary antibody overnight at 4°C. On the next day, sections were washed with PBS, incubated with secondary antibody for 2 h at room temperature, washed again with PBS and incubated for 1 h at room temperature with peanut agglutinin, subsequently washed again and embedded in mounting medium containing DAPI (Vectashield, Vector Laboratories/Biozol, Germany). Whenever needed, the immunostaining was performed on 10 μm thick frozen sections of testis fixed in neutral buffered 4% formalin. The complete list of antibodies and conditions is found in supplemental data (Table S1). Images of stained tissue sections were taken using a fluorescence microscope (Keyence, Bioreva BZ-9000, Germany) or a confocal fluorescence microscope (Leica TCS SPE). For cell counts, at least 100 seminiferous tubuli of 3 mice per group were analyzed using ImageJ software.

### Quantitative realtime PCR

Total RNA was extracted from WT, importin α7^−/−^ and importin α7^ΔIBB/ΔIBB^ testes and FACS sorted germ cells using RNeasy Mini Kits (Qiagen, Hilden, Germany). First-strand DNA synthesis was performed using M-MLV Reverse Transcriptase (Invitrogen, Darmstadt, Germany) and random primers according to the manufacturer’s instructions. Quantitative PCR was performed using Go Taq (Promega, Mannheim, Germany) on an IQ 5 Multicolour Realtime PCR Detection System (Bio-Rad laboratories, München, Germany). Relative gene expression was calculated using the ΔΔCt method with GAPDH as normalizing gene. Primer sequences are listed in supplemented data (Table S2).

### Autoantibody detection

Autoantibodies against sperm proteins were detected as described previously (Meng et al., 2011). Briefly, blots with testicular proteins of two months old WT mice were incubated with a 1:50 dilution of either WT, importin α7^ΔIBB/ΔIBB^ or α7^−/−^ mutant sera overnight at 4°C. Primary antibodies were detected with an IRDye coupled secondary anti-mouse antibody for 1 h at room temperature and detection was performed using the Odyssey Infrared Scanner (LiCor, Bad Homburg, Germany).

### Biotin-labelling of blood-testis barrier

Mice were sacrificed by cervical dislocation and testis were carefully pulled out of the body without extracting them. 50 μl of 1 mM CaCl_2_ containing 10 mg/ml biotin (EZ-Link Sulfo-NHS-LC-Biotin, Pierce, Dallas, USA) were injected with a 0.4 mm needle into one testis. As a control, the second testis was injected with 50 μl of 1 mM CaCl_2_ only. After 30 min of distribution of the injected solution via diffusion the testes were dissected and snap frozen in Tissue-Tek OCT compound (Sakura Finetek, Netherlands). 15 μm cryoslices were cut, mounted on glass slides and fixed with 4% PFA for 15 min. After washing, the sections were incubated with streptavidin-Cy5 directly, coversliped and observed under a fluorescence microscope (Keyence, Bioreva BZ-9000).

### BrdU injection

To analyze proliferation of germ cells, animals received two intraperitoneal injections of bromodeoxyuridine (BrdU; 50 mg/kg body weight dissolved in 0.9% NaCl; Sigma-Aldrich) 2 h apart and were sacrificed 2 h after the second injection.

### RNAsequencing

A total amount of 1 μg RNA per sample was used as input material for the RNA sample preparations. Sequencing libraries were generated using NEBNext® UltraTM RNA Library Prep Kit for Illumina® (NEB, USA) following manufacturer’s recommendations and index codes were added to attribute sequences to each sample. Briefly, mRNA was purified from total RNA using poly-T oligo-attached magnetic beads. Fragmentation was carried out using divalent cations under elevated temperature in NEBNext First Strand Synthesis Reaction Buffer (5X). First strand cDNA was synthesized using random hexamer primer and M-MuLV Reverse Transcriptase (RNase H-). Second strand cDNA synthesis was subsequently performed using DNA Polymerase I and RNase H. Remaining overhangs were converted into blunt ends via exonuclease/polymerase activities. After adenylation of 3’ ends of DNA fragments, NEBNext Adaptor with hairpin loop structure were ligated to prepare for hybridization. In order to select cDNA fragments of preferentially 150~200 bp in length, the library fragments were purified with AMPure XP system (Beckman Coulter, Beverly, USA). Then 3 μl USER Enzyme (NEB, USA) was used with size-selected, adaptor-ligated cDNA at 37 °C for 15 min followed by 5 min at 95 °C before PCR. PCR was performed with Phusion High-Fidelity DNA polymerase, Universal PCR primers and Index (X) Primer. At last, PCR products were purified (AMPure XP system) and library quality was assessed on the Agilent Bioanalyzer 2100 system. Fastq reads were pseudo-aligned to the mm10 genome assembly using kallisto (version 0.46) and transcript read counts were aggregated to Ensembl Gene IDs for further analysis. Differential gene expression analysis was performed via the R library sleuth (Pimentel et al., 2017). Significance and effect sizes of differential gene regulation were calculated from the likelihood ratio and the Wald test, respectively, as implemented in the sleuth package. GO term and pathway enrichment analyses were performed based on the effect size between the WT and knockdown strains using the generally applicable Gen Set Enrichment Analysis (GSEA), GAGE, which determines whether a set of genes is systematically up- or downregulated as a whole (Luo et al., 2009). For gene set definitions, we used the Molecular Signatures Database (MSigDB) from the R msigdf package (Version 7.1) (Liberzon et al., 2015). Gene sets with less than 3 or those with more than 500 members were discarded for statistical robustness and biological interpretation. Putative transcription factor activity from RNA-seq data was assessed per pseudo timepoint against healthy controls using the mouse gene set resource DoRothEA v1, which provides a curated collection of transcription factor and target genes interactions (the regulon) from different sources (Garcia-Alonso et al., 2019). Only interactions with high, likely, and medium confidence (levels A, B, C) were considered. Regulons were statistically evaluated using the R package *viper* (v1.22.0; row-wise t-tests) (Alvarez et al., 2016). Only regulons having at least 15 expressed gene targets were considered.

### Testicular single-cell suspensions

Cells were isolated from 3-months-old WT and α7^ΔIBB/ΔIBB^ male mice according to a protocol of Getun et al. with slight modifications (Getun et al., 2011). Briefly, tunica albuginea was removed, and the seminiferous tubules were fragmented with scissors. The fragments were dissociated in dispase (BD Biosciences) with 10 U/ml DNAse I for 40 min at 37°C. After centrifugation, the pelleted tubules were resuspended in trypLE express enzyme (Life Technologies) with 10 U/ml DNAse I and incubated at 32°C for 20 min. The resulting whole cell suspension was successively washed with Gey’s balanced salt solution (GBSS, Sigma-Aldrich). Then the cell pellet was resuspended in GBSS supplemented with 10% fetal calf serum and 10 U/ml DNAse I. The dissociated testis sample was then passed through a 40 μm GBSS pre-wetted disposable cell strainer. Final staining was performed by adding Hoechst 33342 (5 μg/ml) to the dissociated testis sample and incubating at 32°C for 1 h. Before analysis, propidium iodide (PI at 2 μg/ml) was added to exclude dead cells.

### FACS sorting

FACS sorting was performed according to a slightly modified protocol of Bastos et al. (Bastos et al., 2005). Briefly, the enrichment of pachytene spermatocytes and round spermatids was performed on a FACSAria III cell sorter from BD Biosciences. Live stained testicular cells were excited with a near UV laser (375 nm), the two parameters Hoechst blue (450/40 BP) and Hoechst red (670 LP) were used to identify and sort.

### Extraction of chromatin-bound proteins

Extraction of basic nuclear proteins from mouse testis was performed according to Eckhardt (Eckhardt and Wang-Eckhardt, 2015). Briefly, one testis was homogenized in ice-cold NETN buffer, centrifuged at 12,000 x g for 10 min, resuspended in NETN buffer and centrifuged again. Then, the pellet was resuspended in 0.2 N HCl and incubated overnight at 4°C. After centrifugation at 12,000 x g for 10 min, the supernatant containing basic nuclear proteins was neutralized with 1 M Tris-HCl (pH 8.5) and protein concentration was determined.

### Statistics

Statistical analysis was performed with Prism7 (GraphPad). Results are presented as means ± SEM. Significance was determined by using ANOVA (where 3 groups were compared) or the unpaired two-tailed Student’s t test. For distribution of the genotype after heterozygous mating the binominal test was used. Significance was assumed for p < 0.05 (*, p < 0.05; **, p < 0.01; ***, p < 0.001; ****, p < 0.0001; n.s., not significant).

## Acknowledgements

The authors wish to thank Anne Hahmann, Andrea Rodak and Madeleine Skorna-Nussbeck for technical assistance. We thank Laura Pelz and Fritz Rathjen for providing a BT-IgSF knockout mouse for control of BTB experiments. We also thank Hans-Peter Rahn for his help with the FACS sorting of cells and the Advanced Light Microscopy technology platform of the MDC for their technical support.

## Data Access

The RNAseq data is available through Gene Expression Omnibus (GEO) under the ID GSE160969. Reviewers can access the data at https://www.ncbi.nlm.nih.gov/geo/query/acc.cgi?acc=GSE160969 using the token ‘qxofaykwndszjix’.

## Competing interests

We declare that we have no significant competing financial, professional, or personal interests that might have influenced the performance or presentation of the work described in this manuscript.

## Funding

The work leading to this manuscript was partly supported by the DFG (BA 1374/21-1 and RO 4779/1-2).

---

**Fig. S1** (A) Epididymal sperm count of all homozygous and heterozygous mutant lines including compound heterozygous mice (α7^ΔIBB/−^) compared to WT mice (n=4-6). (B) Western blot analysis of testis protein extracts with N-terminal and C-terminal anti-importin α7 antibodies show, that α7^ΔIBB/−^ testes express both, the full-length and the truncated protein.

**Fig. S2** Principal component analysis of the gene-aggregated expression values as measured in transcipts per million of the 50% most variable genes across all samples.

**Fig. S3** Immunofluorescence for Crem (green) in importin α7^+/+^ and importin α7^ΔIBB/ΔIBB^ testis. For staging of seminiferous tubules, testis paraffin sections were stained with DAPI (blue) and lectin peanut agglutinin (PNA-Al594, red, merge, right panel). Scale bars: 25 μm.

**Table S1** List of antibodies with respective conditions they were used in.

**Table S2** List of primer sequences of target genes.

**Table S3** Differential gene expression in importin α7^ΔIBB/ΔIBB^ and importin α7^−/−^ testes compared to WT and to each other.

**Table S4** List of genes, which are differentially expressed in testis of Rfx2 KO and importin α7^ΔIBB/ΔIBB^ compared to their respective WT controls (P-value cutoff < 0.01, absolute effect size > 0.5).

**Table S5** Gene Set Enrichment Analysis of importin α7^ΔIBB/ΔIBB^ and Rfx2 knockout testes.

